# From single species to communities: microsatellite amplicon sequencing to monitor felids using Feliplex

**DOI:** 10.1101/2024.10.20.619303

**Authors:** Divyashree Rana, Frédéric Boyer, Marta De Barba, Pierre Taberlet, Uma Ramakrishnan

**Affiliations:** National Centre for Biological Sciences, Tata Institute of Fundamental Research, India; Univ. Grenoble Alpes, Univ. Savoie Mont Blanc, CNRS, LECA, Grenoble, France; Biotechnical Faculty, University of Ljubljana, Slovenia; DivjaLabs L.t.d.

**Keywords:** Felids, Genotype-by-sequencing, High-throughput Sequencing, Microsatellite, Multispecies research, Non-invasive

## Abstract

1. The current biodiversity crisis demands a shift from single-species to multispecies approaches in conservation, particularly for rare and endangered species. However, this transition requires tools optimised for multispecies research, which are currently limited.
2. Recent advances in high-throughput sequencing (HTS) technologies and bioinformatics have enabled efficient and robust acquisition of genetic data. Amplicon sequencing approaches, in particular, have demonstrated potential for enhancing non-invasive genetic studies of endangered species, but their application has been mostly limited to single species.
3. To enable multispecies genetic research, we introduce a cost-effective and robust HTS-based amplicon sequencing approach for genotyping multiple species simultaneously, designed for population monitoring, including individual identification and ascertaining patterns of population structure.
4. We developed Feliplex, a multiplex panel of 85 co-amplifying tetranucleotide microsatellite markers for cross-genotyping *Felidae* species, to demonstrate the utility of our approach. Feliplex was validated on known samples from nine Indian felid species across the genera *Panthera, Prionailurus*, and *Felis*. We applied it to invasive (blood and tissue) and non-invasive (hair and faeces) DNA extracts from 173 wild individuals obtaining respectively 70% and 56% multilocus genotyping success rates. The panel accurately identified known population clusters in tigers (*Panthera tigris*, n=19) and revealed hitherto unknown genetic structure in fishing cats (*Prionailurus viverrinus*, n=40).
5. Feliplex’s wide applicability across *Felidae* allows reliable multispecies genotyping from low-quality/quantity samples, while supporting cost-effective genetic studies and conservation monitoring of lesser-known species like small cats. Our approach has a broad applicability and can be adapted to develop similar multispecies panels for closely related species groups.

## Introduction

Conservation biology originated as a crisis discipline (Soulé, 1986). To prioritise efforts and limited resources, certain species were used as proxies for broader co-occurring biodiversity with shared benefits (Caro, 2010). This led to single species focused research andthe development of popular conservation concepts like “umbrella,” “keystone,” “indicator,” or “flagship” species approaches (Simberloff, 1998). However, recent studies have questioned the assumed efficiency of single-species approaches for biodiversity management (Sibarani et al., 2019; Dutta et al., 2023). Consequently, conservation planning has rapidly evolved to incorporate multispecies and ecosystem approaches (Root et al., 2003; Schwenk & Donovan, 2011). However, multispecies assessments require access to detailed and comparable ecological information for multiple species which seldom exists.

Despite a paradigm shift in conservation science towards ecosystem-based approaches, most methodological tools for population studies remain focused on single-species frameworks, limiting data generation for much of biodiversity (Toffelmier et al., 2022). This focus restricts our broader conservation goals, with multispecies assessments often relying on by-catch or opportunistic data from studies aimed at individual species. As knowledge drives attention, single-species assessments tend to concentrate research and funding on charismatic species, diverting attention from lesser-known species (Martin-Lopez et al., 2011). Non-invasive methods such as camera trapping and non-invasive genetic sampling, provide opportunities to study elusive species in their natural habitats (O’Connell, Nichols, & Karanth, 2011; Kelly et al., 2012). While these tools are frequently designed for one or a few target species, they generate data for multiple sympatric species, however with a risk of generating flawed inferences for non-target species (O’Brien, 2008; Hamel et al., 2013). Therefore, for effective multispecies assessment, tools optimised for communities rather than individual species are critical (Toffelmier et al., 2022).

With the advent of technological innovations, genomics has emerged as a crucial tool in understanding evolutionary processes and ecological phenomena, while aiding conservation efforts (van der Valk & Dalén, 2024). Although genetic studies have played an important role in ecological and conservation science for decades, high-throughput technologies have heralded a new genomic era generating significant findings for threatened species (Ouborg et al., 2010; Supple & Shapiro, 2018; Robinson et al., 2019). One important methodological application of genomics is the affordable discovery of nuclear markers for targeted amplification to understand population genetic patterns for non-model organisms (Santana et al., 2009). Amplicon sequencing is particularly effective in elucidating genetic patterns using poor-quality/quantity samples in wild populations (Andrews et al., 2018; Schmidt et al., 2020). This targeted approach can generate high throughput genetic data from a large number of samples by multiplexing a small set of identified informative markers (Campbell et al., 2015). Marker panels optimised for amplicon sequencing of Single Nucleotide Polymorphisms (SNPs) have been applied on various DNA sources, including non-invasive samples, however, their utility is often limited to single species (like Natesh et al., 2019; Burgess et al., 2022). Theoretically, panels targeting multiple species to yield population-level patterns would require cross-amplifying informative markers which showcase polymorphism within and across target species. However, existing multispecies SNP approaches include independent sets of markers which are polymorphic within and across species, hence their applicability being limited to SNPchips with thousands of markers (like SilvalJJunior et al., 2015 and Montanari et al., 2023). These robust species-specific tools create a methodological gap for studying elusive and under-funded species, where access is often limited to poor-quality or non-invasive samples for understanding population trends.

Traditional conservation genetics has relied on microsatellite genotyping for over three decades to understand population biology in the wild (Allendorf, 2017). Contrary to SNPs, microsatellite markers are known for their hypervariable nature and ability to cross-amplify across related species even from degraded and fragmented samples (Moodley & Bruford, 2007). However, these markers have been criticised for being prone to genotyping errors and subjective allele calling, due to their repeated nature (Yuan et al., 2021). Moreover, genotyping based on traditional capillary electrophoresis platforms limits the number of markers that can be analysed in parallel, reducing statistical power and making their use cumbersome. Due to these limitations, their use has been rebuffed, and discrepancies have been highlighted in fine-scale inferences (Supple & Shapiro, 2018; Aylward et al., 2022).

Recent developments in genotyping-by-sequencing of microsatellite markers present a novel opportunity to combine the advantages of hypervariable microsatellites with the parallel processing of high-throughput sequencing (HTS) to efficiently generate replicable data with high statistical power (Bradbury et al., 2018). Although initially applied in human DNA studies (Fordyce et al., 2011), this technique has been successfully tested and optimised for various species, such as grapevines (Kunej et al., 2020), Atlantic cod (Vartia et al., 2016), and brown bears from non-invasively collected samples (De Barba et al., 2017). Moreover, it has proven effective to genotype environmental DNA (eDNA) samples collected from snow tracks of brown bears, wolves, and lynxes (De Barba et al., 2024). The successful application of an HTS approach to microsatellite genotyping in various species suggests the potential for using this method in a multispecies context to optimise robust tools for genotyping several species simultaneously.

Large charismatic species, specifically big cats, receive wide attention globally leading to detailed research and conservation efforts. Genetic and genomic tools have also been used extensively to study them (O’Brien & Johnson, 2005; Li et al., 2016; Prost et al., 2022). However, the *Felidae* family is diverse, comprising 34 species that are phylogenetically closely related and fall into two subfamilies: Pantherinae (big cats) and Felinae (small cats) (Johnson et al., 2006). There is a growing realisation that small carnivores are equally, if not more, important in the changing habitats of the Anthropocene (Marneweck et al., 2024). With the rapid loss of larger predators, smaller, more abundantly distributed, and adaptable small carnivores, like small cats, are critical in maintaining ecological balance. Despite recent attention, critical ecological data on these smaller species is still missing, and the lack of targeted tools hinders research efforts to understand their populations and vulnerabilities.

Here, we introduce Feliplex, a tool that allows the genotyping of multiple species of felids using a set of co-amplifying microsatellite markers sequenced on a HTS platform. Developed on the Illumina MiSeq platform, it generates data for 85 loci and includes a bioinformatic pipeline for genotyping samples from multiple felid species simultaneously. The Feliplex panel was optimised with nine species from three genera to ensure wide applicability across the *Felidae* family. Additionally, we validate the approach for individual identification and population structure using known samples of the tiger (*Panthera tigris*) and unknown samples of the fishing cat (*Prionailurus viverrinus*). Thus, by developing the first-of-its-kind multispecies marker panel, we showcase a new application of genotyping microsatellite markers on an HTS platform.

## Materials and Methods

### Samples and DNA extraction

Samples (n=173) were collected from nine species across three genera: Jungle Cat (JC) *Felis chaus*, Domestic Cat (DC) *Felis catus*, Fishing Cat (FC) *Prionailurus viverrinus*, Leopard Cat (LC) *Prionailurus bengalensis*, Rusty-spotted Cat (RSC) *Prionailurus rubiginosus*, Tiger (TG) *Panthera tigris*, Leopard (LP) *Panthera pardus*, Lion (LN) *Panthera leo*, and Snow Leopard (SL) *Panthera uncia*. The newly designed primer pairs were tested and optimised using multiple DNA sources (tissue, blood, faecal swabs, and hair samples) from these species to ensure wide practical applicability (details in table S4). Additionally, tissue samples from tigers (*Panthera tigris*) and faecal samples from genetically identified fishing cats (*Prionailurus viverrinus*) were used to validate and demonstrate the applicability of the panel for population genetic studies. Faecal samples were collected using sterile swabs in Longmire’s buffer. DNA was extracted from all types of collected samples using the Qiagen DNeasy Blood and Tissue Extraction Kit with manufacturer prescribed protocol.

### Microsatellite discovery and primer design

Primers were designed for de novo discovered candidate microsatellite markers aimed at genotyping low-quality/quantity non-invasive samples using high-throughput sequencing. Based on a previous study (Menotti-Raymond et al., 1999), primers designed on the *Felis catus* genome showed successful amplification across multiple felid species (Beaumont et al, 2001; Biswas et al., 2022; Rana, 2021). Therefore, a domestic cat genome (Accession no. GCF_018350175.1) with minimal missing data (0.9%) was used as the reference genome for primer design. Primer pairs were designed following criteria outlined in De Barba et al. (2017), targeting tetranucleotide repeats with short amplicon sizes (<150 bp) and with similar hybridization temperature (57°C +/-1°C) to allow increased multiplexing from degraded samples.

To ensure multispecies amplification, the de novo designed primers were checked for in-silico amplification using genomes of nine wild felid species from five genera (Table S1). The targeted species with available genomes on the National Centre for Biotechnology Information (NCBI) repository comprise species from four out of the six felid lineages found in India (Johnson et al., 2006). The selected primer pairs were optimised for the Illumina MiSeq platform by adding platform-compatible adapter regions following Natesh et al., 2019 (details provided in SM1).

### Multiplexing conditions and sequencing

To identify the optimal primer pool size for multiplexing, a random subset of primer pairs were tested for singleplex amplification with subsequent gel electrophoresis to determine the common optimal annealing temperature. Markers were then tested in pools of various sizes — 50, 100, and 195 — to ascertain the best pool size. The markers were optimised for a two-step GT-seq library preparation protocol (Campbell et al., 2015) that includes an initial multiplexing reaction followed by an indexing reaction to attach unique barcodes to each sample (as illustrated in Figure 1C). The PCR conditions for the first multiplexing reaction followed the protocol by De Barba et al. (2017) using QIAGEN Multiplex PCR Plus on extracted DNA, while the second indexing reaction followed the protocol by Natesh et al. (2019) (details provided in SM1). The multiplexed and indexed samples were ultimately pooled in an equivolume manner and sequenced on the Illumina MiSeq 300-cycle sequencing platform. Each sample was sequenced three to five times using independent PCR replicates.

**Figure 1.**
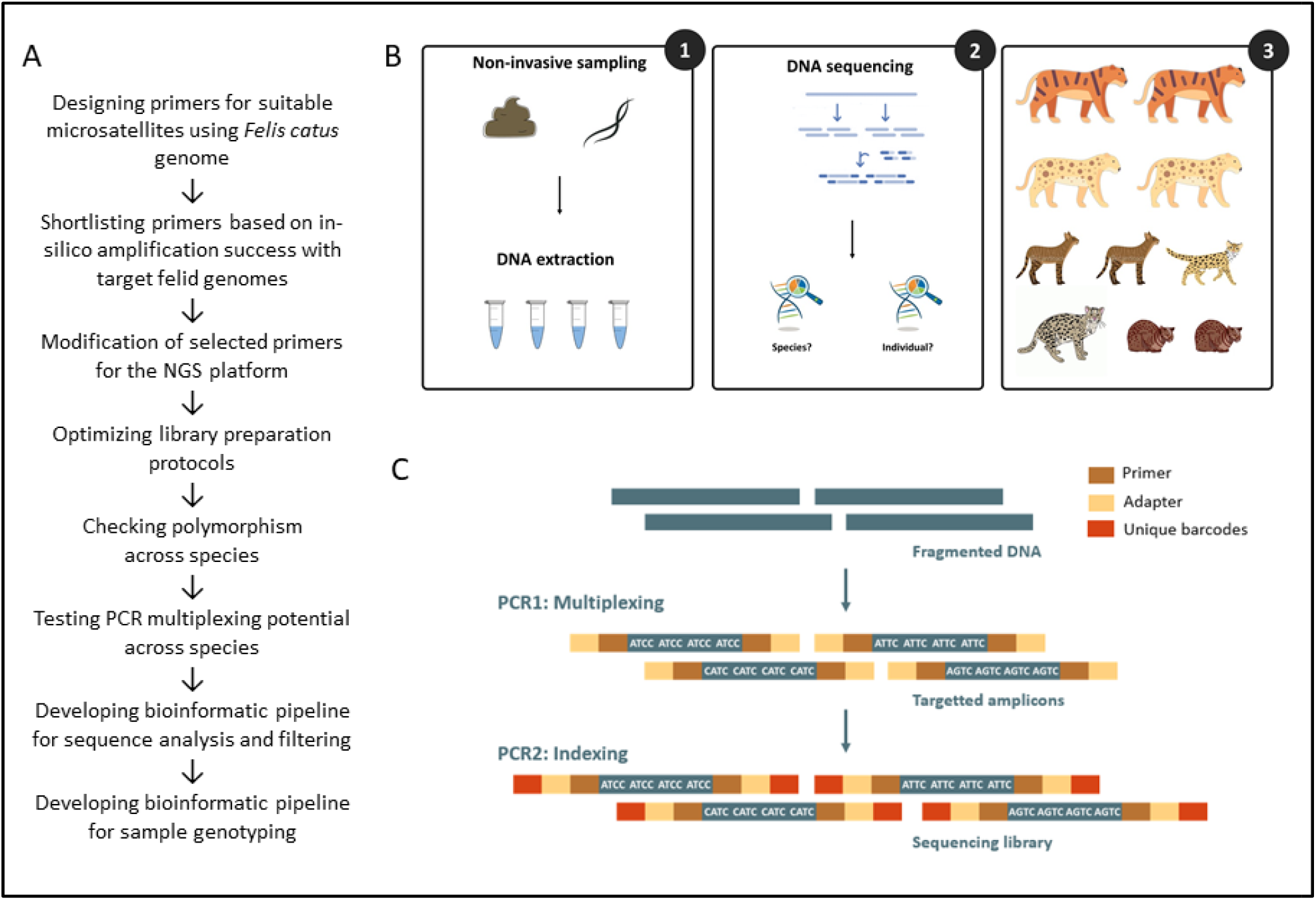
Overview of the panel design and application strategy. A. Development strategy for multispecies Feliplex panel and data generation, B. Schematic describing the application of the designed panel, C. Illustration of the two-step library preparation protocol used in the study.

### Data generation and genotype calling

The raw sequencing reads were trimmed to remove adapters and low-quality sequences (Q>30) using Trimmomatic (Bolger et al., 2014). The R1 and R2 reads were then aligned to generate consensus reads, which were demultiplexed for each primer pair using the *ngsfilter* function in OBITools (Boyer et al., 2016). A read count file was generated from the demultiplexed reads for each marker, containing the sample name, unique amplicon sequences, and their respective counts (read depth). Non-target reads without microsatellite repeats for the locus were excluded from further analysis.

Next, the number of microsatellite repeats in each sequence was counted, creating a table of repeat numbers and their frequencies (read counts) for each sample and marker combination, which was used for allele identification. Alleles were identified based on the relative frequencies of observed repeat sequences, with the most abundant sequences containing the microsatellite motif designated as alleles. Only markers with a depth of >100 per sample were retained. A PCR product was classified as homozygous for a locus if a single allele sequence accounted for over 25% of the total reads, and as heterozygous if two such allele sequences were present. PCR products with more than two alleles, each with a frequency over 25%, were deemed inconclusive. Alleles were identified across all loci for each PCR based on repeat sequence frequencies, resulting in the “genotype dataframe.” This dataframe was filtered and analysed to produce final consensus genotypes with minimal missing data. A consensus genotype at each locus was called when at least three independent replicates produced matching genotypes. PCR replicates with over 90% missing or inconclusive data across loci were discarded before consensus genotypes were called. Genotyping error rates for allelic dropout and false alleles were calculated separately for invasive and non-invasive samples by comparing alleles in each PCR replicate to consensus genotypes, following Broquet & Petit (2004)..

### Marker selection and individual identification

Markers were chosen based on their performance across all tested species, individual genera, and species (TG and FC) using all the samples mentioned above. The “genotype dataframe” was filtered by target taxa, focusing on single or multiple species within a genus as needed. Since microsatellite markers vary in amplification rate (the proportion of PCR replicates yielding at least one allele) and genotyping rate (the proportion of samples yielding a consensus genotype) across species, only markers with an average genotyping rate of 0.4 or higher, and at least two unique alleles across species, were selected. These markers were grouped into two categories, “very high” and “high” efficiency primers, based on read counts during optimization. The selected markers were tested for deviations from Hardy-Weinberg equilibrium (HWE) using *GenAlEx 6*.*5* (Peakall and Smouse, 2006), with significance at each locus determined by a Chi-square test across individuals grouped by geographic populations.

Multilocus consensus genotypes were determined for each sample after establishing consensus genotypes at each locus across replicates. Poor-quality samples and markers, defined as having over 60% missing data across samples or loci, were discarded. Unique individuals were identified using *GenAlEx 6*.*5* (Peakall and Smouse, 2006), based on matching probabilities of multilocus consensus genotypes (Waits et al., 2001). Samples with more than two mismatches, ignoring missing data, were considered recaptures of the same individual. In case of only one or two mismatches, samples were scrutinised to verify if mismatches were true genotypic differences or were caused by genotyping errors. The frequency of null alleles was estimated across loci for all species using the Brookfield (1996) method, implemented through the *PopGenReport* R package (Adamack & Gruber, 2014). Genus-specific genetic clustering of identified individuals was examined using Principal Component Analysis (PCA) and Discriminant Analysis of Principal Components (DAPC) via the *adegenet* R package (Jombart, 2008). The variation explained by the first two principal component axes was used to visualise genetic differentiation between individuals from different species within each genus.

### Testing the panel on known patterns of population structure: Feliplex versus whole genomes for *Panthera tigris*

To validate the utility and accuracy of the developed Feliplex panel to assess population structure, we compared population genetic analyses based on Feliplex-generated genotypes and genomic data. Tissue and blood samples (n=29) from three previously identified population clusters (Natesh et al., 2017) were amplified and genotyped using the Feliplex panel. Simultaneously, whole genomes for the same samples, published in Khan et al. (2021), were used to identify SNPs across the genome. Variant calling and filtering were performed using *bcftools* (Li, 2011) and *vcftools* (Danecek et al., 2011), filtering for biallelic SNPs with a depth of 10-500 and a minimum allele count of 3. Only SNPs present in all samples, with no missing data across samples, were retained.

Both the Feliplex-generated genotypes and the whole-genome SNP data were independently used to assess genetic clustering with Principal Component Analysis (PCA) using the *adegenet* R package (Jombart, 2008) and Bayesian clustering software STRUCTURE version 2.3.4 (Pritchard, Stephens, and Donnelly, 2000). STRUCTURE was run under default settings, using ancestry models with correlated frequencies. The number of population clusters (K) was tested from 1 to 5, with 10 independent runs for each K. The burn-in period and MCMC repetitions were set to 100,000 and 1,000,000, respectively. The optimal K was selected based on the maximum log likelihood of the posterior probability of the data at each K, as well as delta K measures (Evanno et al., 2005). Additionally, Puechmaille’s method (Puechmaille, 2016) was applied to account for uneven sampling across populations, using the StructureSelector software (Li & Lui, 2018). Lastly, pairwise Fst values between identified clusters were calculated using the *adegenet* R package. This approach validated the Feliplex genotypes by comparing them to genotype data derived from whole-genome sequences to assess population structure patterns.

### Application of the panel to assess unknown patterns of population structure: Feliplex with faecal samples of *Prionailurus viverrinus*

The utility of the panel to ascertain population-level patterns from non-invasive samples was tested using faecal samples (n=75) collected during field surveys across the fishing cat range in India. These samples were gathered from four states: Andhra Pradesh (AP), Orissa (OD), West Bengal (WB) on the Eastern coast, and Uttar Pradesh (UP) in the Northern floodplains, as shown in Figure 4D. The samples were confirmed to be from *Prionailurus viverrinus* based on sequencing results using an independent felid-specific 16s mitochondrial primer (Mukherjee et al. 2016), as part of a previous study (Rana, 2021). Samples were amplified and genotyped using the Feliplex panel as described above. Without prior genetic information, samples were grouped into putative populations based on their geographical locations and proximity. Genetic clustering and distance between the geographic clusters were assessed using PCA, STRUCTURE (for k = 1 to 10), and pairwise Fst, as explained above.

## Results

### Microsatellite discovery and selection

A total of 1292 markers were identified using the *Felis catus* reference genome. In-silico amplification with the target species produced 195 primer pairs with a single target amplicon, allowing up to 2 mismatches per binding site, but none at the 3’ end. The average pooled library size, inferred from TapeStation, ranged between 180 bp and 220 bp due to variable repeats across species. Consequently, the libraries were targeted for 120 bp read lengths on the 2×150bp Illumina MiSeq platform.

The 195 primer pairs were tested for amplification across multiple species, with pools of varying sizes. Primer efficiency significantly dropped for pools larger than 100. However, better performance was achieved by amplifying samples in two pools of “very-high” and “high” efficiency primers, compared to a single primer pool during the initial library preparation step. After filtering markers based on primer amplification efficiency and genotyping rates across species, 85 markers were retained. Different optimal “very-high” and “high” efficiency primer pools were identified for each target genus (details in SM3). Among selected markers, 10-15% showed significant deviation from HWE (Table S9). Genotyping error rates were low, with average allelic dropout rates of 0.011 and 0.025, and false allele rates of 0.008 and 0.004, for invasive and non-invasive samples, respectively (locus specific error rates are provided in Table S10).

### Multispecies discrimination and individual identification

The panel successfully genotyped 103 samples across 45 loci from nine felid species, each forming distinct clusters (Figure 2A). Group-wise PID and PIDsibs were low (PID: 8.20E-04 to 2.50E-31, PIDsibs: 2.80E-02 to 9.30E-14, SM3), providing high confidence in individual identification. Asiatic lions (LN) had the highest PID values, whereas domestic cats (DC) had the lowest values. Additionally, known unique LN individuals mismatched at less than two loci for multiple samples, reflecting their low genetic variation and isolated population (Bertola et al., 2015). Selected markers were polymorphic, with polymorphism ranging from 1 to 8 alleles/locus per species (Figure 2B). DC, FC, and TG exhibited the highest polymorphism within their genera. Null allele frequencies were below 0.2 on average for the majority of the loci across species (Figure S3), in concordance to multiple studies (Dakin & Avise, 2004).

**Figure 2.**
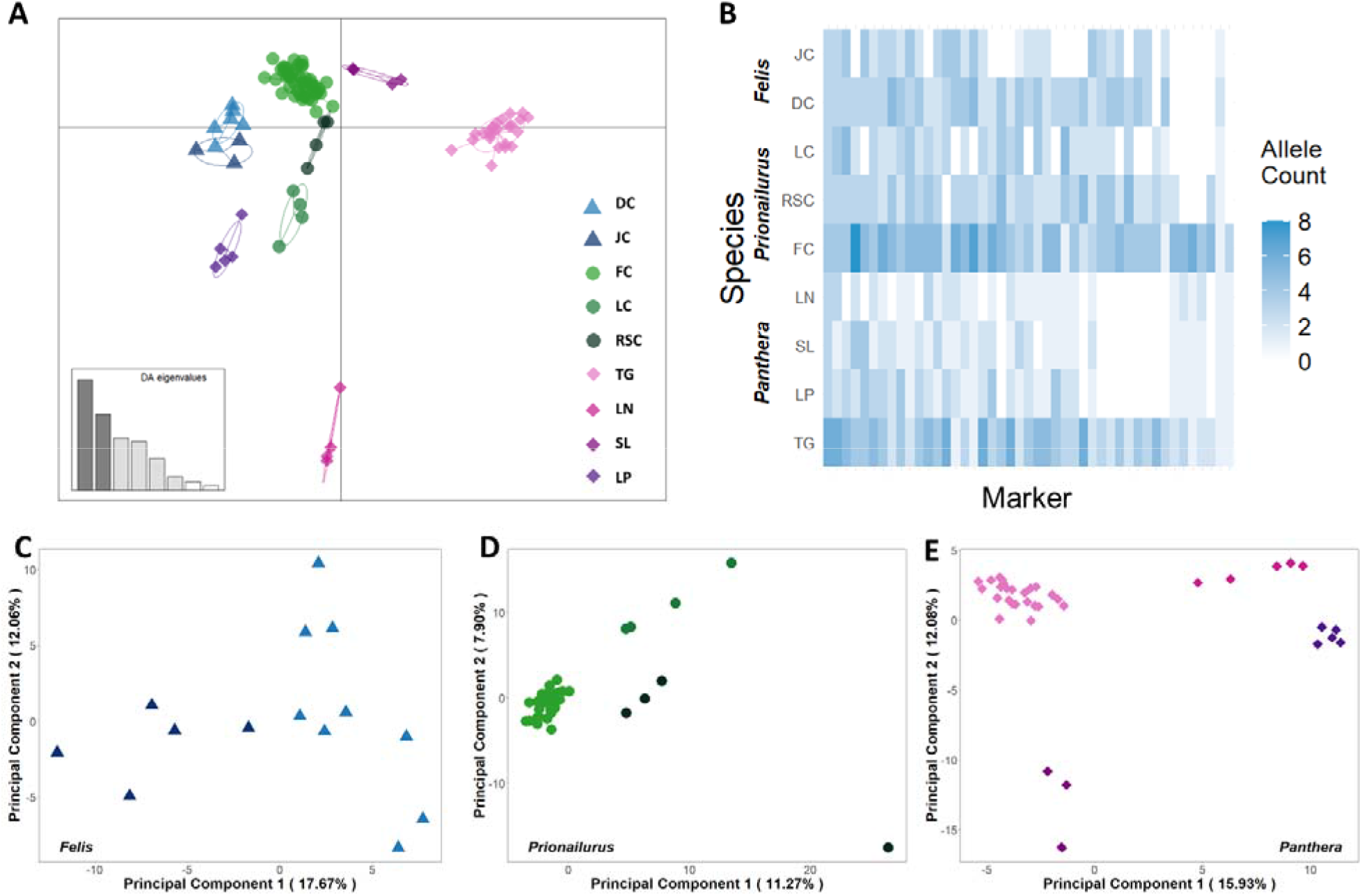
Application of Feliplex panel for seven felid species. A. Discriminant Principal Analysis showing segregation of samples based on species. B. Heatmap illustrating polymorphism for the cross-amplifying 31 markers successfully genotyped for different species. C-E. Principal Component Analysis using genus-specific markers to demonstrate species-wise segregation for C. *Felis* (Domestic Cat DC and Jungle Cat JC), D. *Prionailurus* (Fishing Cat FC, Leopard Cat LC, and Rusty-spotted Cat RSC), and E. *Panthera* (Tiger TG, Leopard LP, Lion LN, Snow Leopard SL).

Primer efficiency varied across species, requiring different primer pools for different groups. Multilocus genotyping success rates varied from 53% to 73% across species (see Table S5). When filtering markers for individual identification by genus, 40 loci for Felis (Figure 2C), 46 for Prionailurus (Figure 2D), and 38 for Panthera (Figure 2E) were retained. One locus for Prionailurus and four for Panthera were monomorphic. Species within each genus clustered independently separating distinctly on the PCA axes., with the first two PC axes explaining 29.73% of variation within Felis, 19.17% within Prionailurus, and 27.47% within Panthera (Figures 2C-2E).

### Testing the panel on known patterns of population structure: Feliplex versus whole genomes for *Panthera tigris*

Using blood and tissue samples from 19 successfully genotyped individuals across three known genetic clusters—Central India (CI, n=6), North West (NW, n=8), and Southern India (SI, n=5)— we performed a comparative analysis. The samples were analysed using 55 loci from the Feliplex panel, and 28,627 biallelic SNPs from published whole genome data. Out of the 55 loci, 15 were monomorphic, leaving 40 informative markers for further analysis.

Three unique genetic clusters were identified using the STRUCTURE algorithm, with distinct populations forming independent genetic clusters. This pattern was evident in both SNPs from whole genomes (Figure 3-A1) and amplicon data using Feliplex markers (Figure 3-B1). PCA analysis also accurately revealed three genetic clusters, with the first two principal components (PCs) explaining 32.79% and 39.12% of the variation in genomic and amplicon data, respectively (Figures 3-A2 and 3-B2). Although both datasets explained a similar percentage of variation, the whole genome data provided a clearer resolution. Pairwise Fst values between clusters ranged from 0.16 to 0.29 based on whole genome data, with the highest differentiation between NW and SI populations. Feliplex-based pairwise Fst values were slightly higher, ranging from 0.28 to 0.31, but both datasets showed similar population differentiation patterns.

**Figure 3.**
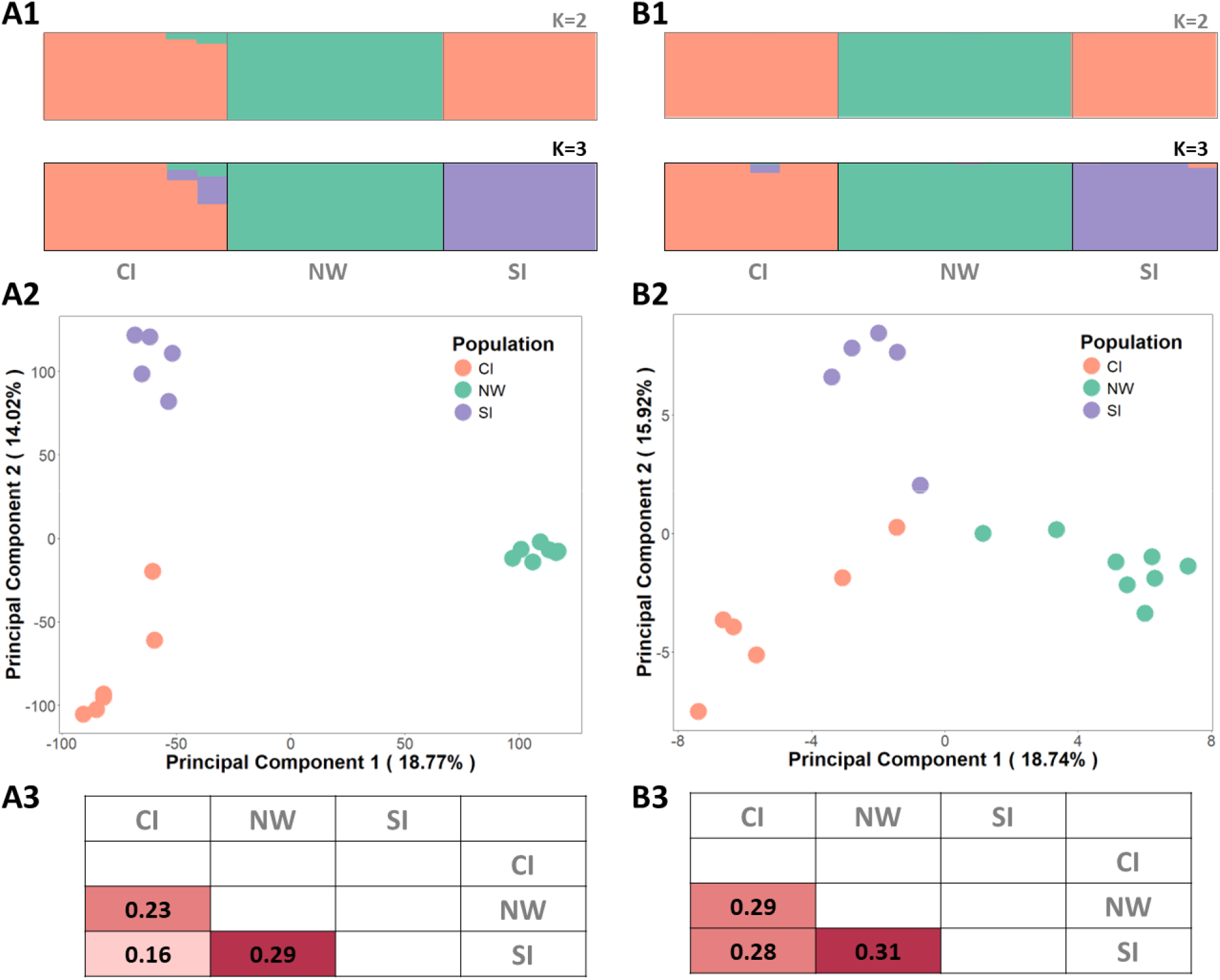
Validation of the Feliplex panel using known tiger high-quality samples. Population genetic analysi (1. STRUCTURE, 2. Principal Component Analysis, 3. Pairwise Fst) conducted using A. Whole genome data and B. Amplicon sequencing data using Feliplex. The optimal K (K=3) is highlighted in figures A1 and A2.

### Application of the panel to assess unknown patterns of population structure: Feliplex with faecal samples of *Prionailurus viverrinus*

Using Feliplex, consensus genotypes were generated for 40 samples at 36 tetranucleotide repeat loci. Based on multilocus matching, six samples were identified as recaptures. Discarding the recaptures and four monomorphic loci, a 32-locus genotype for each of 34 unique individuals from across the country was used for population genetic analysis. The PCA analysis revealed the genetic structuring of the samples based on geographical proximit (Figure 4A). Bayesian clustering using STRUCTURE suggested presence and admixture between three probable genetic clusters (K=3), based on the log-likelihood approach as well as the Puechmaille’s method. Samples from the southernmost sampled population on the Eastern Coast (AP), were found to be genetically most distinct. Other samples from OD and WB clustered together forming a unique genetic cluster, based on the identified optimal K clustering. On the other hand, samples from northern floodplains in UP showcased the presence of two genetic clusters with admixture with populations in WB and OD. (Figure 4A and 4B). Genetic distance between sampling locations was found to be high, ranging from 0.05-0.31, as calculated by pairwise Fst. The highest pairwise Fst value (0.31) was recorded between WB and AN, whereas the lowest value (0.05) was recorded between geographically distant UP and OD populations. The OD population exhibited least differentiation with other populations, with high similarity with WB as well as UP population.

**Figure 4.**
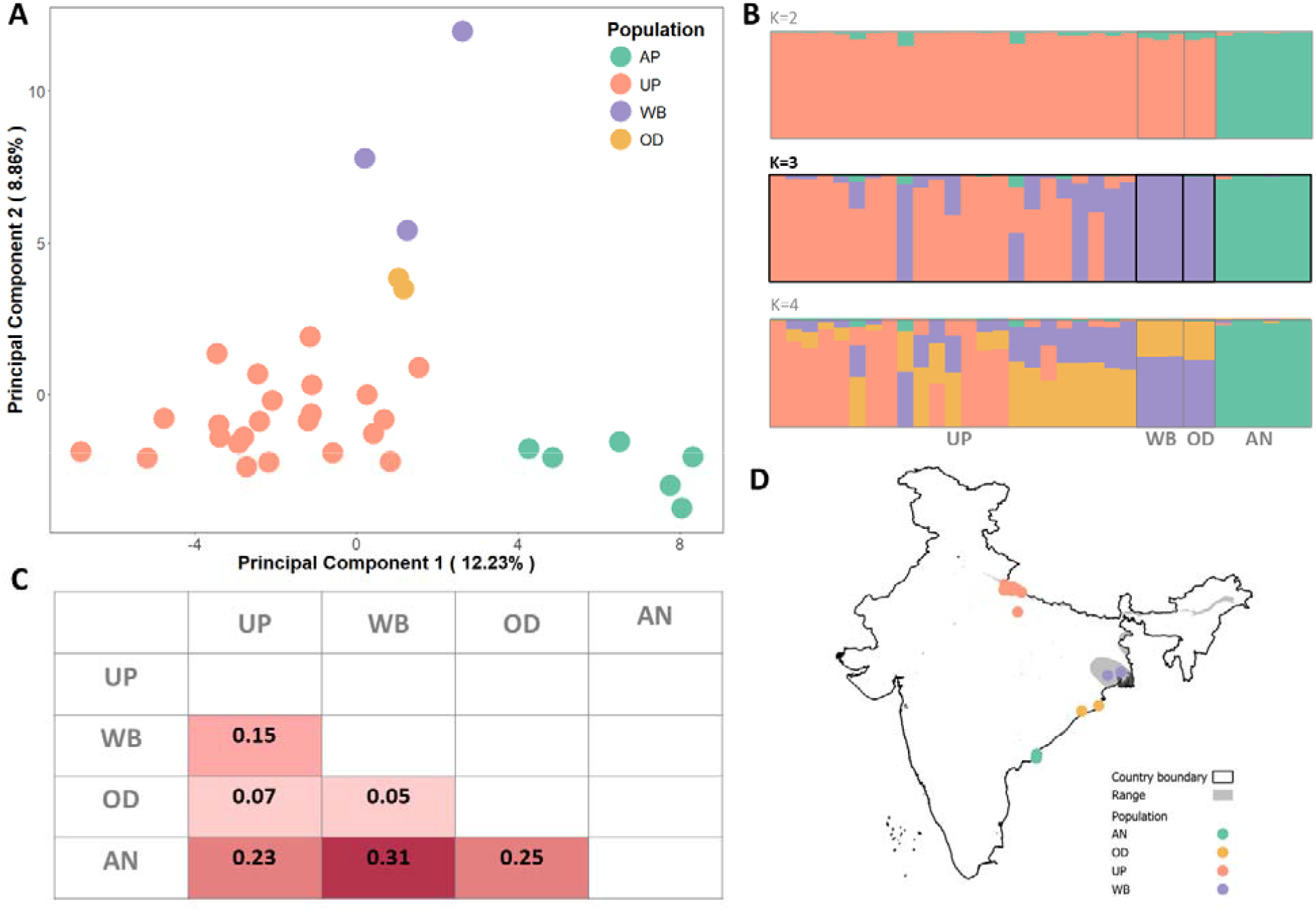
Assessment of population structure for fishing cat faecal samples. Population genetic analysis. A. Principal Component Analysis, B. STRUCTURE, C. Pairwise Fst, and D. Sampling locations. The optimal K (K=3) is highlighted in figure B.

## Discussion

In the new era of conservation biology, multispecies-centric research requires robust data for multiple sympatric species, often in contrast to the available species-specific tools. Here, utilising genotype-by-sequencing of cross-species hypervariable microsatellite markers, we present an optimised tool capable of genotyping non-invasive samples from multiple wild felid species. We discovered and optimised a set of 85 co-amplifying markers which demonstrate positive cross-amplification across the *Felidae* family, focusing on genera found in India. Using a two-step GT-seq library preparation protocol, the Feliplex marker panel was optimised for high-throughput sequencing platform followed by a bioinformatic pipeline for generating consensus genotypes. With a set of over 30 co-amplifying Feliplex markers for nine tested species, the accuracy and utility of the markers for population genetic analyses was demonstrated using known tiger and unknown fishing cat samples. Overall, our results demonstrate the utility of our approach to successfully identify individuals and reveal population-level patterns for multiple species across *Felidae* simultaneously.

### Wide applicability across *Felidae*

Leveraging the phylogenetic relatedness of felids, markers originally designed for domestic cats (Menotti-Raymond et al., 1999) have been extensively used to genotype wild felid species. However, our initial efforts to utilise pre-existing cross-amplifying dinucleotide markers in a HTS multiplexing panel revealed low multiplexing capacities. To address this limitation, we focused on developing a broader set of tetranucleotide co-amplifying markers, optimised for efficient sequencing on an HTS platform.

Amplicon sequencing has demonstrated its reliability in generating robust genetic data, even from degraded samples (Andrews et al., 2018). In a novel approach to support multispecies research, the Feliplex markers were designed for amplicon sequencing across multiple species. India, recognized as a global priority region for felid conservation and home to the world’s highest diversity of felids (Dickman et al., 2015), provided an ideal testing ground for this panel. The Feliplex markers successfully produced accurate individual identification for nine species across three diverse genera in India, illustrating their broad applicability across taxa. Furthermore, the potential relevance of this panel for Neotropical felids, a closely related clade, merits further exploration.

Compared to big cats, small cats are elusive and under-studied, often receiving limited research funding. In several regions, small cat taxonomy is still being revised with the discovery of hybrids and new species (Trigo et al., 2013; De Oliveira et al., 2024). This study aims to provide a standardised tool for generating reliable and reproducible genetic data for lesser-known small cat species, which often suffer from insufficient funding and a lack of long-term population monitoring. One compelling example of the Feliplex markers’ utility is their application to the study of the fishing cat, a threatened wetland specialist with a fragmented distribution and largely unknown patterns of population structure and connectivity (Rana et al., 2022). Our findings indicate (a) high genetic diversity among fishing cats in the northern floodplains of UP, (b) gene flow between populations in the northern floodplains and the northern east coast regions (WB and OD), and (c) isolation of the southernmost populations in AP. Notably, fishing cat populations in OD appear connected to those in both WB and UP, suggesting possible connectivity through central India, a hypothesis that warrants further investigation. Sporadic photographic evidence of fishing cats from central India over the past decade further supports the need for additional studies on their presence in this region (Talegaonkar et al., 2018; Dutta et al., 2021). The multispecies approach facilitated by the Feliplex panel enhances the efficiency of data generation and analysis, particularly in habitats where multiple felids coexist sympatrically.

### Efficiency in understanding population structure

Our results underscored the multispecies applicability of the Feliplex panel for accurate individual identification, but we also sought to evaluate the accuracy and limitations of this approach in determining population structure. Tigers, being the most studied felids in India with well-documented population structure from high-resolution data (Natesh et al., 2017), provided an ideal reference for comparison. We compared data generated using the Feliplex panel to available whole-genome data. The patterns observed were consistent with published findings, validating the panel’s ability to detect fine-scale population structure. Both the tens of thousands of SNPs from whole genomes and the ∼45 Feliplex microsatellites revealed the presence of three known genetic clusters with little to no admixture between them (Figure 3). While the first two PC axes explained similar variation (∼35%) in both datasets, the genetic patterns were more finely resolved with the high-resolution genomic data, as expected. Nevertheless, the Feliplex panel successfully identified unique individuals across all populations, including those in the highly related and inbred northwestern population (Khan et al., 2021). Thus, we conclude that the Feliplex markers are sufficiently powerful for individual identification and for reliably determining population-level parameters. However, for highly related or inbred populations, detecting fine-scale genetic patterns may benefit from genomic approaches with greater statistical power. Our data revealed similar locus-specific frequency of null alleles as reported in literature, however impact of null alleles should be further evaluated in field application studies with larger sample sizes. Ultimately, the utility of the Feliplex panel should be considered in relation to the specific research question and population characteristics. It is especially valuable for exploring populations with no prior genetic information.

### Utility and cost-effectiveness of the approach

In addition to the research question, the choice of method is often constrained by practical factors such as the availability of technology or dedicated funding. Traditionally, microsatellites have been analysed using fragment analysis via capillary electrophoresis, which is limited to a small number of samples (typically 96) and markers (usually fewer than 10 per sample) per run. This method is both cost- and labour-intensive, offering low power and resolution to address complex questions (Guichoux et al., 2011). In contrast, genotyping-by-sequencing with Feliplex markers allows simultaneous data generation from multiple loci and samples, overcoming these limitations.

The quality of DNA extracted from different sample types varies greatly. Tissue samples typically yield intact, high-concentration DNA, whereas faecal samples often produce low-concentration, degraded (i.e., fragmented) DNA. Therefore, a method’s utility for understanding population-level parameters using low-quality, non-invasive samples depends on its ability to generate reliable data. Sequencing approaches, from amplicon sequencing to whole genome sequencing, have variable success rates with faecal samples (White et al., 2019). For instance, while double digest Restriction-Enzyme Associated Digestion (ddRAD) can generate genomic data from thousands of SNPs, this method struggles with low-quantity faecal DNA (Andrews et al., 2018). Additionally, ddRAD requires large amounts of template DNA, often leading to significant sample loss during processing, with fewer than 20% of collected samples making it through analysis (Tyagi et al., 2024). As a result, methods like ddRAD may not be feasible for research on elusive carnivores. In contrast, amplicon sequencing of microsatellites has been successfully applied to both non-invasive and eDNA samples (De Barba et al., 2017; De Barba et al., 2024; Barbian et al., 2018). In our study, genotyping success—the proportion of samples with multilocus consensus genotypes—was on average 56% for non-invasive samples and 70% for invasive samples, with a considerably higher number of loci analysed compared to typical microsatellite studies. Replicating samples by performing several independent PCR reactions from a single sample is crucial for generating reliable consensus genotypes, particularly for non-invasive samples (Taberlet et al., 1996). We replicated each sample three to five times and discarded those with high levels of missing data. If needed, higher replication levels could be efficiently achieved with a HTS approach, which would increase overall genotyping success (De Barba et al., 2017).

Cost is another critical factor when choosing a method for studying wild populations. In India, for instance, ddRAD processing costs around 50 USD per sample, while whole genome sequencing at 20x depth costs approximately 500 USD per sample. In contrast, the estimated cost of processing a sample with Feliplex amplicon sequencing is about 15 USD, including DNA extraction and sequencing, though excluding labour and computational expenses. While the current approach utilises the Illumina MiSeq platform, it could be adapted to higher-throughput platforms like NovaSeq 6000, which would reduce the per-sample cost by at least five times. Lastly, traditional marker discovery for single species is effective but expensive, ideally requiring available genomes from all target populations (Natesh et al., 2019; Delord et al., 2018). Feliplex provides a cost-effective solution for felid research in resource-limited settings, even in the absence of extensive genomic data.

### Caveats and recommendations

The phylogenetic relatedness among felid species ensures that Feliplex primers co-amplify across the Felidae family. However, due to documented variations in primer efficiency between species, optimising Feliplex primers for species-specific use would enhance genotyping success. Alternatively, for studies requiring fewer markers, researchers can select only the “very-high” efficiency primers for genotyping. As mentioned earlier, replication of sample amplification is essential for reliable genotyping. In this study, we applied a moderate level of sample replication (three to five PCR replicates per sample). In cases where prior knowledge suggests a low probability of generating consensus genotypes, a more targeted approach could involve screening samples for amplification success and only repeating those with higher success rates.

Incorporating species and sex identification markers at low concentrations into the primer pool could expand the application of amplicon panels (as demonstrated by De Barba et al., 2017). However, adding a mitochondrial primer to the Feliplex pool resulted in an unequal distribution of reads, given the higher copy number of mitochondrial DNA, hence, it was dropped. Loci with private alleles for certain species or populations could be used in assignment tests, but this would require building a more extensive dataset including samples from multiple species and populations. In this study, efforts were made to include samples from several populations across India. A few monomorphic loci were detected with private alleles in different species, but their amplification was inconsistent, making them unsuitable as species identification markers in the multiplex. Future standardisation, supported by broader datasets, could resolve this issue. Despite genus-specific optimization, access to raw reads, allele sequences, and the bioinformatics pipeline used for sequence data analysis facilitates comparison and reproducibility of results. This addresses a major criticism against the use of traditional microsatellites for genotyping.

## Conclusion

The primary goal of this study was to develop co-amplifying nuclear markers capable of generating individual-level genetic data for multiple felid species. Historically, microsatellite markers have been crucial in understanding the genetics of many threatened and elusive species, leveraging the power of co-amplifying markers and hypervariable loci. The Feliplex panel offers a cost-effective approach by harnessing multispecies markers using a HTS platform, while addressing technical criticisms of microsatellite genotyping. Our panel has demonstrated broad applicability across the *Felidae* family, enabling individual identification and the detection of population-level parameters from degraded samples. Future research with additional felid species and expanded applications, such as investigating hybridization in small cats, will further validate and refine this approach, advancing its use in conservation research. Moreover, the workflow we used to develop Feliplex can be used to develop similar multispecies panels for other closely related taxa. Lastly, we believe this approach would specifically aid in assisting research on species with limited prior genomic information.

## Supporting information

Summplementary Material

## Acknowledgements

DR is supported by the TIFR-NCBS graduate program. We acknowledge the Panthera Foundation Small Cat Action Fund awarded to DR for their funding support in field testing and optimising the Feliplex panel. Our gratitude extends to the National Centre for Biological Sciences for their institutional support as facilitated by UR, specifically the Next-Generation Genomics Facility (NGGF, Bangalore Life Science Cluster, BLiSC) for their assistance in data generation. The NCBS data cluster, supported under project no. 12-R&D-TFR-5.04-0900 by the Department of Atomic Energy, Government of India, was indispensable. We thank Vinay Sagar, Divyajyoti Ganguly, and Denkant Singha for providing samples to test the panel’s applicability. Lastly, DR is grateful to Imran and Nova for their crucial inputs in formulating this study.

## Data Accessibility

Genotype data, data processing scripts and supplementary data are available on Dryad.

