## Supplementary material for "From single species to communities: microsatellite amplicon sequencing to monitor felids using Feliplex": Summplementary Material

### SM1. Microsatellite discovery and Primer design

**Table S1. Details of the available genomes used in marker discovery and selection**

| **Species** | **Common Name** | **Lineage** | **Data type** | **Accession No.** | **Application** |
| --- | --- | --- | --- | --- | --- |
| *Felis catus* | Domestic Cat | Domestic Cat Lineage | Reference genome | GCF_018350175.1 | Reference genome for marker discovery |
| *Felis chaus* | Jungle Cat | Domestic Cat Lineage | Reference genome | GCA_019924945.1 | In-silico amplification check |
| *Felis silvestris* | Wildcat | Domestic Cat Lineage | SRA Reads | ERX10012593 |  |
| *Prionailurus viverrinus* | Fishing Cat | Leopard Cat Lineage | Reference genome | GCA_022837055.1 |  |
| *Prionailurus bengalensis* | Leopard Cat | Leopard Cat Lineage | Reference genome | GCF_016509475.1 |  |
| *Prionailurus rubiginosus* | Rusty-spotted Cat | Leopard Cat Lineage | SRA Reads | SRX3213487 |  |
| *Caracal caracal* | Caracal | Caracal Lineage | Reference genome | GCA_016801355.1 |  |
| *Panthera leo* | Lion | Panthera Lineage | Reference genome | GCF_018350215.1 |  |
| *Panthera pardus* | Leopard | Panthera Lineage | Reference genome | GCF_024362965.1 |  |
| *Panthera tigris* | Tiger | Panthera Lineage | Reference genome | GCF_018350195.1 |  |
| *Panthera uncia* | Snow Leopard | Panthera Lineage | Reference genome | GCF_023721935.1 |  |
| *Neofelis nebulosa* | Clouded Leopard | Panthera Lineage | Reference genome | GCA_028018385.1 |  |

Designed primers were modified with the following overhang compatible with the unique barcodes, 5’-CGACAGGTTCAGAGTTCTACAGTCCGACGATC-3’ (overhang with forward primer) and 5’-GTGACTGGAGTTCAGACGTGTGCTCTTCCGATCT-3’ (overhang with reverse primer).

### SM2. Library preparation protocol and optimization

The primer pairs were divided into “very-high” and “high”-efficiency pools. The first multiplexing PCR was conducted separately for both pools with each sample, and the products were combined for the second indexing PCR.

The first PCR was conducted in 10uL reactions which included 5uL Qiagen Multiplex PCR Plus MM, 1uL of primer pool, and 4uL of DNA extract. The first multiplexing PCR conditions following De Barba et al., 2017 included denaturation at 95°C for 2 minutes followed by 45 cycles of 30 seconds at 95°C, 90 seconds at 57°C, 60 seconds at 72°C, and a final elongation step of 10 minutes at 72°C. DNA concentration of a few random samples was checked using a Qubit Fluorometer. If the concentration ranged between 5-15ng/uL, indexing PCR was performed. If the sample concentration was more than the range, the PCR products from the first reaction were diluted 10 times with 4uL of PCR1 product and 36uL of nuclease-free water. The PCR1 products from both primer pools were pooled in an equal volume of 5uL each.

The second PCR was conducted in 15uL reactions which included 6uL Qiagen Multiplex PCR MM, 3uL of i5 (forward) indexes, 3uL of i7 (reverse) indexes, and 3uL of PCR1 product. The index combinations have to be unique to be pooled in a common library which could be demultiplexed post sequencing. The second indexing PCR conditions following Natesh et al., 2019 included initial denaturation at 95°C for 15 minutes followed by 12 cycles of 30 seconds at 95°C, 30 seconds at 62°C, 30 seconds at 72°C, and a final elongation step of 5 min at 72°C. The indexed PCR products were pooled in an equal volume of 5uL each. The pooled library was size-selected twice to retain fragments between 100-400 bp using AMPureXP beads. Pooled DNA extracts from ten samples from each species were used to optimise laboratory protocols and check the multispecies efficiency of the Feliplex markers.

### SM3. Marker and sample selection

| **Table S2. Genotyping success rate of markers across species** | | | | | | | | | | | |
| --- | --- | --- | --- | --- | --- | --- | --- | --- | --- | --- | --- |
|  | **Marker** | **Felis** | | **Prionailurus** | | | **Panthera** | | | | **average** |
|  |  | **JC** | **DC** | **LC** | **RSC** | **FC** | **LN** | **SL** | **LP** | **TG** |  |
| **All species** | **FPX106** | **1** | **0.67** | **0.75** | **0.75** | **0.91** | **NA** | **0.75** | **1** | **1** | **0.85** |
|  | **FPX11** | **1** | **0.8** | **1** | **1** | **0.2** | **0.45** | **1** | **1** | **0.89** | **0.82** |
|  | **FPX127** | **0.8** | **0.89** | **NA** | **1** | **0.89** | **1** | **1** | **1** | **0.98** | **0.94** |
|  | **FPX13** | **0.9** | **1** | **1** | **0.75** | **0.99** | **NA** | **NA** | **NA** | **0.63** | **0.88** |
|  | **FPX159** | **1** | **1** | **1** | **1** | **0.9** | **0.9** | **1** | **0.8** | **0.87** | **0.94** |
|  | **FPX163** | **0.33** | **0.33** | **NA** | **0.25** | **0.33** | **0.9** | **1** | **1** | **0.89** | **0.63** |
|  | **FPX172** | **NA** | **NA** | **NA** | **NA** | **0.68** | **1** | **1** | **1** | **1** | **0.94** |
|  | **FPX176** | **0.33** | **0.5** | **NA** | **0.25** | **0.98** | **NA** | **NA** | **NA** | **0.54** | **0.52** |
|  | **FPX180** | **1** | **1** | **1** | **1** | **0.65** | **0.68** | **1** | **1** | **0.96** | **0.92** |
|  | **FPX185** | **0.26** | **1** | **0.75** | **0.75** | **1** | **0.9** | **NA** | **1** | **0.65** | **0.79** |
|  | **FPX186** | **0.67** | **0.5** | **NA** | **0.25** | **0.91** | **NA** | **NA** | **NA** | **0.54** | **0.57** |
|  | **FPX19** | **NA** | **NA** | **NA** | **0.25** | **0.67** | **1** | **1** | **1** | **0.96** | **0.81** |
|  | **FPX195** | **NA** | **0.83** | **1** | **1** | **0.83** | **NA** | **NA** | **NA** | **0.33** | **0.8** |
|  | **FPX197** | **NA** | **1** | **NA** | **0.25** | **0.69** | **0.78** | **1** | **1** | **0.74** | **0.78** |
|  | **FPX204** | **NA** | **NA** | **NA** | **NA** | **0.74** | **0.22** | **1** | **1** | **0.74** | **0.74** |
|  | **FPX206** | **1** | **0.8** | **1** | **1** | **0.32** | **0.68** | **1** | **1** | **0.94** | **0.86** |
|  | **FPX207** | **0.9** | **0.95** | **0.5** | **0.5** | **0.8** | **NA** | **0.75** | **1** | **0.74** | **0.77** |
|  | **FPX215** | **NA** | **0.33** | **0.75** | **0.75** | **0.99** | **1** | **1** | **1** | **1** | **0.85** |
|  | **FPX219** | **NA** | **0.17** | **1** | **0.75** | **0.94** | **NA** | **NA** | **NA** | **1** | **0.77** |
|  | **FPX227** | **0.33** | **0.17** | **0.25** | **0.25** | **0.53** | **1** | **1** | **1** | **0.76** | **0.59** |
|  | **FPX228** | **NA** | **0.17** | **0.25** | **0.5** | **0.91** | **1** | **1** | **1** | **0.87** | **0.71** |
|  | **FPX232** | **NA** | **0.17** | **0.5** | **0.25** | **0.89** | **1** | **1** | **1** | **0.87** | **0.71** |
|  | **FPX238** | **NA** | **0.17** | **0.25** | **0.25** | **0.29** | **1** | **0.75** | **1** | **0.96** | **0.58** |
|  | **FPX244** | **0.67** | **0.33** | **1** | **1** | **0.5** | **1** | **0.75** | **1** | **0.78** | **0.78** |
|  | **FPX258** | **0.8** | **0.7** | **NA** | **0.25** | **0.58** | **1** | **1** | **1** | **0.98** | **0.79** |
|  | **FPX259** | **NA** | **NA** | **NA** | **NA** | **0.71** | **1** | **1** | **1** | **0.98** | **0.94** |
|  | **FPX268** | **1** | **1** | **1** | **0.75** | **0.78** | **NA** | **0.5** | **1** | **0.76** | **0.85** |
|  | **FPX278** | **0.8** | **1** | **1** | **1** | **0.96** | **NA** | **1** | **1** | **0.92** | **0.96** |
|  | **FPX280** | **0.9** | **0.75** | **1** | **0.5** | **0.95** | **1** | **1** | **1** | **0.8** | **0.88** |
|  | **FPX32** | **NA** | **0.67** | **0.75** | **0.75** | **0.83** | **NA** | **NA** | **NA** | **0.58** | **0.72** |
|  | **FPX33** | **NA** | **NA** | **0.25** | **0.25** | **0.88** | **NA** | **NA** | **NA** | **0.62** | **0.5** |
|  | **FPX38** | **NA** | **NA** | **NA** | **0.25** | **0.86** | **NA** | **NA** | **NA** | **0.54** | **0.55** |
|  | **FPX39** | **0.33** | **0.17** | **0.75** | **0.25** | **0.96** | **0.9** | **1** | **1** | **0.92** | **0.7** |
|  | **FPX4** | **NA** | **0.17** | **0.75** | **0.5** | **0.88** | **0.45** | **1** | **1** | **0.96** | **0.71** |
|  | **FPX49** | **0.9** | **0.95** | **0.75** | **1** | **0.35** | **1** | **0.75** | **1** | **0.89** | **0.84** |
|  | **FPX55** | **0.67** | **0.17** | **0.5** | **0.25** | **0.88** | **0.9** | **1** | **1** | **0.98** | **0.7** |
|  | **FPX69** | **0.67** | **0.5** | **0.75** | **NA** | **0.68** | **1** | **1** | **1** | **0.94** | **0.82** |
|  | **FPX7** | **NA** | **0.17** | **1** | **0.5** | **0.84** | **1** | **1** | **1** | **0.85** | **0.8** |
|  | **FPX72** | **NA** | **NA** | **NA** | **NA** | **0.47** | **0.22** | **1** | **1** | **0.91** | **0.72** |
|  | **FPX76** | **0.9** | **0.89** | **1** | **1** | **1** | **1** | **1** | **1** | **0.96** | **0.97** |
|  | **FPX80** | **0.8** | **0.62** | **1** | **0.75** | **1** | **NA** | **NA** | **NA** | **0.04** | **0.7** |
|  | **FPX83** | **0.9** | **0.89** | **NA** | **1** | **0.63** | **NA** | **1** | **1** | **0.82** | **0.89** |
|  | **FPX85** | **0.8** | **0.89** | **NA** | **1** | **0.82** | **NA** | **NA** | **NA** | **0.68** | **0.84** |
|  | **FPX91** | **0.8** | **0.8** | **0.5** | **0.75** | **0.81** | **NA** | **NA** | **NA** | **0.5** | **0.69** |
|  | **FPX99** | **1** | **0.95** | **1** | **1** | **0.12** | **0.9** | **0.5** | **1** | **0.82** | **0.81** |
| **Felis** | **FPX105** | **0.8** | **0.89** | **NA** | **NA** | **NA** | **NA** | **NA** | **NA** | **NA** | **0.84** |
|  | **FPX155** | **NA** | **0.78** | **NA** | **NA** | **NA** | **NA** | **NA** | **NA** | **NA** | **0.78** |
|  | **FPX16** | **0.6** | **0.56** | **NA** | **NA** | **NA** | **NA** | **NA** | **NA** | **NA** | **0.58** |
|  | **FPX165** | **1** | **0.67** | **NA** | **NA** | **NA** | **NA** | **NA** | **NA** | **NA** | **0.84** |
|  | **FPX168** | **1** | **0.78** | **NA** | **NA** | **NA** | **NA** | **NA** | **NA** | **NA** | **0.89** |
|  | **FPX169** | **0.6** | **0.67** | **NA** | **NA** | **NA** | **NA** | **NA** | **NA** | **NA** | **0.64** |
|  | **FPX178** | **1** | **1** | **NA** | **NA** | **NA** | **NA** | **NA** | **NA** | **NA** | **1** |
|  | **FPX183** | **1** | **1** | **NA** | **NA** | **NA** | **NA** | **NA** | **NA** | **NA** | **1** |
|  | **FPX184** | **0.4** | **0.67** | **NA** | **NA** | **NA** | **1** | **1** | **1** | **0.92** | **0.83** |
|  | **FPX2** | **1** | **0.89** | **NA** | **NA** | **NA** | **NA** | **NA** | **NA** | **NA** | **0.95** |
|  | **FPX20** | **0.8** | **0.67** | **NA** | **NA** | **NA** | **NA** | **NA** | **NA** | **NA** | **0.74** |
|  | **FPX257** | **0.6** | **0.56** | **NA** | **NA** | **NA** | **NA** | **NA** | **NA** | **NA** | **0.58** |
|  | **FPX262** | **1** | **1** | **1** | **1** | **0.35** | **NA** | **NA** | **NA** | **NA** | **0.87** |
|  | **FPX270** | **0.6** | **0.89** | **NA** | **NA** | **NA** | **NA** | **NA** | **NA** | **NA** | **0.74** |
|  | **FPX74** | **1** | **0.11** | **NA** | **NA** | **NA** | **NA** | **NA** | **NA** | **NA** | **0.56** |
|  | **FPX75** | **0.8** | **1** | **NA** | **NA** | **NA** | **NA** | **NA** | **NA** | **0.55** | **0.78** |
|  | **FPX78** | **NA** | **0.89** | **NA** | **NA** | **NA** | **NA** | **NA** | **NA** | **NA** | **0.89** |
|  | **FPX87** | **1** | **0.78** | **1** | **1** | **0.37** | **NA** | **NA** | **NA** | **NA** | **0.83** |
|  | **FPX94** | **0.8** | **0.44** | **NA** | **NA** | **NA** | **NA** | **NA** | **NA** | **NA** | **0.62** |
| **Prionailurus** | **FPX10** | **NA** | **NA** | **0.25** | **0.25** | **0.93** | **NA** | **NA** | **NA** | **NA** | **0.48** |
|  | **FPX115** | **NA** | **NA** | **NA** | **0.25** | **0.63** | **NA** | **NA** | **NA** | **0.65** | **0.51** |
|  | **FPX139** | **NA** | **NA** | **NA** | **0.25** | **0.67** | **NA** | **NA** | **NA** | **NA** | **0.46** |
|  | **FPX150** | **NA** | **NA** | **1** | **1** | **0.57** | **NA** | **NA** | **NA** | **NA** | **0.86** |
|  | **FPX154** | **NA** | **NA** | **NA** | **NA** | **0.7** | **NA** | **NA** | **NA** | **NA** | **0.7** |
|  | **FPX24** | **NA** | **NA** | **1** | **0.25** | **0.59** | **NA** | **NA** | **NA** | **NA** | **0.61** |
|  | **FPX45** | **NA** | **NA** | **0.5** | **1** | **0.65** | **NA** | **NA** | **NA** | **NA** | **0.72** |
| **Panthera** | **FPX125** | **NA** | **NA** | **NA** | **NA** | **NA** | **0.8** | **1** | **1** | **0.85** | **0.91** |
|  | **FPX141** | **NA** | **NA** | **NA** | **NA** | **NA** | **0.6** | **NA** | **NA** | **0.58** | **0.59** |
|  | **FPX142** | **NA** | **NA** | **NA** | **NA** | **NA** | **0.6** | **1** | **1** | **0.96** | **0.89** |
|  | **FPX147** | **NA** | **NA** | **NA** | **NA** | **NA** | **1** | **1** | **1** | **0.67** | **0.92** |
|  | **FPX152** | **NA** | **NA** | **NA** | **NA** | **NA** | **0.8** | **1** | **1** | **0.74** | **0.88** |
|  | **FPX189** | **NA** | **NA** | **NA** | **NA** | **NA** | **NA** | **1** | **NA** | **0.82** | **0.91** |
|  | **FPX210** | **NA** | **NA** | **NA** | **NA** | **NA** | **0.8** | **1** | **1** | **0.82** | **0.91** |
|  | **FPX220** | **NA** | **NA** | **NA** | **NA** | **NA** | **0.6** | **1** | **1** | **0.98** | **0.89** |
|  | **FPX229** | **NA** | **NA** | **NA** | **NA** | **NA** | **1** | **1** | **1** | **0.76** | **0.94** |
|  | **FPX273** | **NA** | **NA** | **NA** | **NA** | **NA** | **1** | **1** | **1** | **0.78** | **0.95** |
|  | **FPX274** | **NA** | **NA** | **NA** | **NA** | **NA** | **NA** | **1** | **1** | **0.96** | **0.98** |
|  | **FPX3** | **NA** | **NA** | **NA** | **NA** | **NA** | **NA** | **1** | **1** | **0.82** | **0.94** |
|  | **FPX54** | **NA** | **NA** | **NA** | **NA** | **NA** | **1** | **1** | **NA** | **0.68** | **0.89** |
|  | **FPX84** | **NA** | **NA** | **NA** | **NA** | **NA** | **NA** | **NA** | **NA** | **0.6** | **0.6** |
| **Primer Count** | | **45** | **57** | **37** | **48** | **54** | **40** | **46** | **45** | **62** |  |

| **Table S3. Amplification rate of markers across species** | | | | | | | | | | | |
| --- | --- | --- | --- | --- | --- | --- | --- | --- | --- | --- | --- |
|  | **Marker** | **Felis** | | **Prionailurus** | | | **Panthera** | | | | **average** |
|  |  | **JC** | **DC** | **LC** | **RSC** | **FC** | **LN** | **SL** | **LP** | **TG** |  |
| **All species** | **FPX106** | **0.9** | **0.96** | **0.79** | **1** | **0.8** | **NA** | **1** | **0.83** | **0.96** | **0.9** |
|  | **FPX11** | **0.9** | **0.3** | **0.64** | **0.6** | **0.09** | **0.13** | **0.92** | **0.92** | **0.76** | **0.58** |
|  | **FPX127** | **0.4** | **0.52** | **NA** | **0.6** | **0.59** | **1** | **1** | **0.83** | **0.87** | **0.73** |
|  | **FPX13** | **0.8** | **0.78** | **0.93** | **0.9** | **0.74** | **NA** | **NA** | **NA** | **0.56** | **0.78** |
|  | **FPX159** | **1** | **0.96** | **0.86** | **0.8** | **0.64** | **0.47** | **1** | **0.92** | **0.92** | **0.84** |
|  | **FPX163** | **0.1** | **0.09** | **NA** | **0.2** | **0.22** | **0.6** | **1** | **0.83** | **0.8** | **0.48** |
|  | **FPX172** | **NA** | **NA** | **NA** | **NA** | **0.44** | **1** | **1** | **1** | **1** | **0.89** |
|  | **FPX176** | **0.1** | **0.17** | **NA** | **0.2** | **0.8** | **NA** | **NA** | **NA** | **0.37** | **0.33** |
|  | **FPX180** | **0.9** | **0.52** | **0.64** | **0.6** | **0.33** | **0.4** | **1** | **0.92** | **0.85** | **0.68** |
|  | **FPX185** | **0.3** | **1** | **0.86** | **1** | **0.76** | **0.67** | **NA** | **0.92** | **0.67** | **0.77** |
|  | **FPX186** | **0.2** | **0.17** | **NA** | **0.2** | **0.81** | **NA** | **NA** | **NA** | **0.4** | **0.36** |
|  | **FPX19** | **NA** | **NA** | **NA** | **0.1** | **0.52** | **1** | **1** | **0.92** | **0.83** | **0.73** |
|  | **FPX195** | **NA** | **0.3** | **0.64** | **0.6** | **0.68** | **NA** | **NA** | **NA** | **0.24** | **0.49** |
|  | **FPX197** | **NA** | **0.57** | **NA** | **0.1** | **0.39** | **0.6** | **1** | **0.83** | **0.6** | **0.58** |
|  | **FPX204** | **NA** | **NA** | **NA** | **NA** | **0.55** | **0.07** | **1** | **0.83** | **0.63** | **0.62** |
|  | **FPX206** | **0.7** | **0.35** | **0.57** | **0.6** | **0.18** | **0.6** | **0.92** | **0.83** | **0.89** | **0.63** |
|  | **FPX207** | **0.6** | **0.7** | **0.14** | **0.2** | **0.69** | **NA** | **0.58** | **0.83** | **0.59** | **0.54** |
|  | **FPX215** | **NA** | **0.09** | **0.21** | **0.4** | **0.55** | **1** | **1** | **0.92** | **0.87** | **0.63** |
|  | **FPX219** | **NA** | **0.04** | **0.29** | **0.4** | **0.58** | **NA** | **NA** | **NA** | **0.92** | **0.45** |
|  | **FPX227** | **0.1** | **0.04** | **0.14** | **0.1** | **0.35** | **1** | **1** | **0.83** | **0.65** | **0.47** |
|  | **FPX228** | **NA** | **0.04** | **0.07** | **0.2** | **0.94** | **1** | **1** | **1** | **0.72** | **0.62** |
|  | **FPX232** | **NA** | **0.04** | **0.14** | **0.1** | **0.75** | **1** | **1** | **0.92** | **0.89** | **0.6** |
|  | **FPX238** | **NA** | **0.04** | **0.07** | **0.1** | **0.13** | **1** | **1** | **0.92** | **0.84** | **0.51** |
|  | **FPX244** | **0.3** | **0.09** | **0.57** | **0.6** | **0.19** | **1** | **0.75** | **0.92** | **0.67** | **0.57** |
|  | **FPX258** | **0.3** | **0.43** | **NA** | **0.1** | **0.48** | **1** | **1** | **0.83** | **0.93** | **0.63** |
|  | **FPX259** | **NA** | **NA** | **NA** | **NA** | **0.5** | **1** | **1** | **0.92** | **0.92** | **0.87** |
|  | **FPX268** | **0.9** | **0.91** | **0.79** | **0.4** | **0.65** | **NA** | **0.42** | **0.75** | **0.63** | **0.68** |
|  | **FPX278** | **0.3** | **0.83** | **0.93** | **0.9** | **0.82** | **NA** | **1** | **0.92** | **0.95** | **0.83** |
|  | **FPX280** | **0.5** | **0.61** | **0.64** | **0.3** | **0.56** | **1** | **1** | **0.92** | **0.73** | **0.7** |
|  | **FPX32** | **NA** | **0.26** | **0.36** | **0.3** | **0.67** | **NA** | **NA** | **NA** | **0.48** | **0.41** |
|  | **FPX33** | **NA** | **NA** | **0.07** | **0.1** | **0.58** | **NA** | **NA** | **NA** | **0.39** | **0.28** |
|  | **FPX38** | **NA** | **NA** | **NA** | **0.1** | **0.69** | **NA** | **NA** | **NA** | **0.32** | **0.37** |
|  | **FPX39** | **0.1** | **0.04** | **0.21** | **0.1** | **0.72** | **0.67** | **0.92** | **1** | **0.91** | **0.52** |
|  | **FPX4** | **NA** | **0.04** | **0.21** | **0.2** | **0.61** | **0.13** | **1** | **0.92** | **0.89** | **0.5** |
|  | **FPX49** | **0.8** | **0.74** | **0.64** | **0.5** | **0.18** | **0.6** | **1** | **0.83** | **0.91** | **0.69** |
|  | **FPX55** | **0.2** | **0.04** | **0.14** | **0.1** | **0.76** | **0.47** | **1** | **0.83** | **0.91** | **0.49** |
|  | **FPX69** | **0.2** | **0.13** | **0.21** | **NA** | **0.33** | **1** | **1** | **0.92** | **0.96** | **0.59** |
|  | **FPX7** | **NA** | **0.04** | **0.57** | **0.2** | **0.65** | **1** | **1** | **1** | **0.75** | **0.65** |
|  | **FPX72** | **NA** | **NA** | **NA** | **NA** | **0.3** | **0.07** | **1** | **0.92** | **0.8** | **0.62** |
|  | **FPX76** | **0.7** | **0.65** | **0.71** | **0.8** | **0.68** | **1** | **1** | **0.92** | **0.84** | **0.81** |
|  | **FPX80** | **0.3** | **0.35** | **0.36** | **0.3** | **0.67** | **NA** | **NA** | **0.08** | **0.01** | **0.3** |
|  | **FPX83** | **0.6** | **0.65** | **NA** | **0.6** | **0.4** | **NA** | **0.92** | **0.75** | **0.79** | **0.67** |
|  | **FPX85** | **0.3** | **0.52** | **NA** | **0.6** | **0.5** | **NA** | **NA** | **0.08** | **0.57** | **0.43** |
|  | **FPX91** | **0.4** | **0.35** | **0.29** | **0.4** | **0.74** | **NA** | **NA** | **NA** | **0.41** | **0.43** |
|  | **FPX99** | **0.8** | **0.43** | **0.64** | **0.6** | **0.05** | **0.6** | **1** | **0.83** | **0.72** | **0.63** |
| **Felis** | **FPX105** | **0.7** | **0.57** | **NA** | **NA** | **NA** | **NA** | **NA** | **NA** | **NA** | **0.64** |
|  | **FPX155** | **NA** | **0.57** | **NA** | **NA** | **NA** | **NA** | **NA** | **NA** | **NA** | **0.57** |
|  | **FPX16** | **0.3** | **0.35** | **NA** | **NA** | **NA** | **NA** | **NA** | **NA** | **NA** | **0.32** |
|  | **FPX165** | **0.7** | **0.3** | **NA** | **NA** | **NA** | **NA** | **NA** | **NA** | **NA** | **0.5** |
|  | **FPX168** | **0.8** | **0.35** | **NA** | **NA** | **NA** | **NA** | **NA** | **NA** | **NA** | **0.58** |
|  | **FPX169** | **0.3** | **0.39** | **NA** | **NA** | **NA** | **NA** | **NA** | **NA** | **NA** | **0.34** |
|  | **FPX178** | **0.9** | **0.57** | **NA** | **NA** | **NA** | **NA** | **NA** | **NA** | **NA** | **0.74** |
|  | **FPX183** | **0.9** | **0.48** | **NA** | **NA** | **NA** | **NA** | **NA** | **NA** | **NA** | **0.69** |
|  | **FPX184** | **0.2** | **0.48** | **NA** | **NA** | **NA** | **1** | **1** | **0.92** | **0.92** | **0.75** |
|  | **FPX2** | **0.8** | **0.43** | **NA** | **NA** | **NA** | **NA** | **NA** | **NA** | **NA** | **0.62** |
|  | **FPX20** | **0.8** | **0.26** | **NA** | **NA** | **NA** | **NA** | **NA** | **NA** | **NA** | **0.53** |
|  | **FPX257** | **0.3** | **0.43** | **NA** | **NA** | **NA** | **NA** | **NA** | **NA** | **NA** | **0.36** |
|  | **FPX262** | **0.9** | **0.57** | **0.64** | **0.6** | **0.14** | **NA** | **NA** | **NA** | **NA** | **0.57** |
|  | **FPX270** | **0.3** | **0.43** | **NA** | **NA** | **NA** | **NA** | **NA** | **NA** | **NA** | **0.36** |
|  | **FPX74** | **0.9** | **0.13** | **NA** | **NA** | **NA** | **NA** | **NA** | **NA** | **NA** | **0.52** |
|  | **FPX75** | **0.8** | **0.83** | **NA** | **NA** | **NA** | **NA** | **NA** | **NA** | **0.59** | **0.74** |
|  | **FPX78** | **NA** | **0.57** | **NA** | **NA** | **NA** | **NA** | **NA** | **NA** | **NA** | **0.57** |
|  | **FPX87** | **0.9** | **0.39** | **0.57** | **0.5** | **0.2** | **NA** | **NA** | **NA** | **NA** | **0.51** |
|  | **FPX94** | **0.6** | **0.17** | **NA** | **NA** | **NA** | **NA** | **NA** | **NA** | **NA** | **0.38** |
| **Prionailurus** | **FPX10** | **NA** | **NA** | **0.07** | **0.1** | **0.65** | **NA** | **NA** | **NA** | **NA** | **0.27** |
|  | **FPX115** | **NA** | **NA** | **NA** | **0.2** | **0.39** | **NA** | **NA** | **NA** | **0.55** | **0.38** |
|  | **FPX139** | **NA** | **NA** | **NA** | **0.1** | **0.45** | **NA** | **NA** | **NA** | **NA** | **0.28** |
|  | **FPX150** | **NA** | **NA** | **0.57** | **0.6** | **0.27** | **NA** | **NA** | **NA** | **NA** | **0.48** |
|  | **FPX154** | **NA** | **NA** | **NA** | **NA** | **0.51** | **NA** | **NA** | **NA** | **NA** | **0.51** |
|  | **FPX24** | **NA** | **NA** | **0.71** | **0.4** | **0.41** | **NA** | **NA** | **NA** | **NA** | **0.51** |
|  | **FPX45** | **NA** | **NA** | **0.14** | **0.5** | **0.39** | **NA** | **NA** | **NA** | **NA** | **0.34** |
| **Panthera** | **FPX125** | **NA** | **NA** | **NA** | **NA** | **NA** | **0.6** | **0.92** | **0.83** | **0.77** | **0.78** |
|  | **FPX141** | **NA** | **NA** | **NA** | **NA** | **NA** | **0.53** | **NA** | **NA** | **0.52** | **0.52** |
|  | **FPX142** | **NA** | **NA** | **NA** | **NA** | **NA** | **0.6** | **0.92** | **1** | **0.89** | **0.85** |
|  | **FPX147** | **NA** | **NA** | **NA** | **NA** | **NA** | **1** | **1** | **0.92** | **0.52** | **0.86** |
|  | **FPX152** | **NA** | **NA** | **NA** | **NA** | **NA** | **0.6** | **0.83** | **0.83** | **0.64** | **0.72** |
|  | **FPX189** | **NA** | **NA** | **NA** | **NA** | **NA** | **NA** | **1** | **NA** | **0.72** | **0.86** |
|  | **FPX210** | **NA** | **NA** | **NA** | **NA** | **NA** | **0.6** | **1** | **0.83** | **0.79** | **0.8** |
|  | **FPX220** | **NA** | **NA** | **NA** | **NA** | **NA** | **0.2** | **0.92** | **0.83** | **0.83** | **0.7** |
|  | **FPX229** | **NA** | **NA** | **NA** | **NA** | **NA** | **0.93** | **0.92** | **0.92** | **0.65** | **0.86** |
|  | **FPX273** | **NA** | **NA** | **NA** | **NA** | **NA** | **1** | **1** | **0.92** | **0.6** | **0.88** |
|  | **FPX274** | **NA** | **NA** | **NA** | **NA** | **NA** | **NA** | **1** | **0.92** | **0.87** | **0.93** |
|  | **FPX3** | **NA** | **NA** | **NA** | **NA** | **NA** | **NA** | **1** | **0.83** | **0.67** | **0.83** |
|  | **FPX54** | **NA** | **NA** | **NA** | **NA** | **NA** | **1** | **1** | **NA** | **0.52** | **0.84** |
|  | **FPX84** | **NA** | **NA** | **NA** | **NA** | **NA** | **NA** | **NA** | **NA** | **0.52** | **0.52** |
| **Primer Count** | | **45** | **57** | **37** | **48** | **54** | **40** | **46** | **47** | **62** |  |

| **Table S4. Sample details** | | | | |
| --- | --- | --- | --- | --- |
| **S.No.** | **Sample Name** | **Species** | **Genus** | **Type** |
| **1** | **112-001-IFX-1i** | **DC** | **Felis** | **Faecal** |
| **2** | **17-002-IFX-1i** | **DC** | **Felis** | **Faecal** |
| **3** | **28-002-IFX-2ii** | **DC** | **Felis** | **Faecal** |
| **4** | **3-008-IFX-1i** | **DC** | **Felis** | **Faecal** |
| **5** | **39-004-IFX-1i** | **DC** | **Felis** | **Faecal** |
| **6** | **NOVA** | **DC** | **Felis** | **Blood** |
| **7** | **198-002-IFX-1i** | **DC** | **Felis** | **Faecal** |
| **8** | **4-003-IFX-1i** | **DC** | **Felis** | **Faecal** |
| **9** | **4-008-JKL-1i** | **DC** | **Felis** | **Faecal** |
| **10** | **201920-DDW14** | **JC** | **Felis** | **Faecal** |
| **11** | **ANM22F40** | **JC** | **Felis** | **Faecal** |
| **12** | **STR18-33** | **JC** | **Felis** | **Faecal** |
| **13** | **202223-DDW44** | **JC** | **Felis** | **Faecal** |
| **14** | **202223-DDW6** | **JC** | **Felis** | **Faecal** |
| **15** | **H10** | **LN** | **Panthera** | **Hair** |
| **16** | **H19** | **LN** | **Panthera** | **Hair** |
| **17** | **H36** | **LN** | **Panthera** | **Hair** |
| **18** | **H38** | **LN** | **Panthera** | **Hair** |
| **19** | **H42** | **LN** | **Panthera** | **Hair** |
| **20** | **LC001** | **LP** | **Panthera** | **Blood** |
| **21** | **LC002** | **LP** | **Panthera** | **Blood** |
| **22** | **LC003** | **LP** | **Panthera** | **Blood** |
| **23** | **LF001** | **LP** | **Panthera** | **Blood** |
| **24** | **MZ** | **LP** | **Panthera** | **Blood** |
| **25** | **EAR** | **SL** | **Panthera** | **Tissue** |
| **26** | **SL2** | **SL** | **Panthera** | **Blood** |
| **27** | **SL3** | **SL** | **Panthera** | **Blood** |
| **28** | **SL34A** | **SL** | **Panthera** | **Blood** |
| **29** | **CI15** | **TG** | **Panthera** | **Tissue** |
| **30** | **CI18** | **TG** | **Panthera** | **Tissue** |
| **31** | **CI19** | **TG** | **Panthera** | **Tissue** |
| **32** | **CI2** | **TG** | **Panthera** | **Tissue** |
| **33** | **CI4** | **TG** | **Panthera** | **Blood** |
| **34** | **CI7** | **TG** | **Panthera** | **Blood** |
| **35** | **NW10** | **TG** | **Panthera** | **Blood** |
| **36** | **NW16** | **TG** | **Panthera** | **Blood** |
| **37** | **NW19** | **TG** | **Panthera** | **Blood** |
| **38** | **NW20** | **TG** | **Panthera** | **Blood** |
| **39** | **NW24** | **TG** | **Panthera** | **Blood** |
| **40** | **NW5** | **TG** | **Panthera** | **Blood** |
| **41** | **NW55** | **TG** | **Panthera** | **Blood** |
| **42** | **NW6** | **TG** | **Panthera** | **Blood** |
| **43** | **SI1** | **TG** | **Panthera** | **Blood** |
| **44** | **SI11** | **TG** | **Panthera** | **Blood** |
| **45** | **SI3** | **TG** | **Panthera** | **Blood** |
| **46** | **SI6** | **TG** | **Panthera** | **Blood** |
| **47** | **SI8** | **TG** | **Panthera** | **Blood** |
| **48** | **SI9** | **TG** | **Panthera** | **Blood** |
| **49** | **T57** | **TG** | **Panthera** | **Blood** |
| **50** | **201920-COR1** | **FC** | **Prionailurus** | **Faecal** |
| **51** | **201920-DDW1** | **FC** | **Prionailurus** | **Faecal** |
| **52** | **201920-DDW15** | **FC** | **Prionailurus** | **Faecal** |
| **53** | **201920-DDW18** | **FC** | **Prionailurus** | **Faecal** |
| **54** | **201920-DDW19** | **FC** | **Prionailurus** | **Faecal** |
| **55** | **201920-DDW20** | **FC** | **Prionailurus** | **Faecal** |
| **56** | **201920-DDW21** | **FC** | **Prionailurus** | **Faecal** |
| **57** | **201920-DDW24** | **FC** | **Prionailurus** | **Faecal** |
| **58** | **201920-DDW25** | **FC** | **Prionailurus** | **Faecal** |
| **59** | **201920-DDW28** | **FC** | **Prionailurus** | **Faecal** |
| **60** | **201920-DDW3** | **FC** | **Prionailurus** | **Faecal** |
| **61** | **201920-DDW4** | **FC** | **Prionailurus** | **Faecal** |
| **62** | **201920-DDW5** | **FC** | **Prionailurus** | **Faecal** |
| **63** | **201920-DDW6** | **FC** | **Prionailurus** | **Faecal** |
| **64** | **201920-DDW7** | **FC** | **Prionailurus** | **Faecal** |
| **65** | **201920-DDW8** | **FC** | **Prionailurus** | **Faecal** |
| **66** | **201920-DLR2** | **FC** | **Prionailurus** | **Faecal** |
| **67** | **201920-DLR3** | **FC** | **Prionailurus** | **Faecal** |
| **68** | **201920-JZO1** | **FC** | **Prionailurus** | **Faecal** |
| **69** | **201920-KAT1** | **FC** | **Prionailurus** | **Faecal** |
| **70** | **201920-KAT16** | **FC** | **Prionailurus** | **Faecal** |
| **71** | **201920-KAT3** | **FC** | **Prionailurus** | **Faecal** |
| **72** | **201920-KAT4** | **FC** | **Prionailurus** | **Faecal** |
| **73** | **201920-KAT5** | **FC** | **Prionailurus** | **Faecal** |
| **74** | **201920-KIS1** | **FC** | **Prionailurus** | **Faecal** |
| **75** | **201920-KIS11** | **FC** | **Prionailurus** | **Faecal** |
| **76** | **201920-KIS6** | **FC** | **Prionailurus** | **Faecal** |
| **77** | **201920-LZO3** | **FC** | **Prionailurus** | **Faecal** |
| **78** | **201920-POR1** | **FC** | **Prionailurus** | **Faecal** |
| **79** | **201920-POR2** | **FC** | **Prionailurus** | **Faecal** |
| **80** | **201920-POR3** | **FC** | **Prionailurus** | **Faecal** |
| **81** | **201920-POR4** | **FC** | **Prionailurus** | **Faecal** |
| **82** | **201920-POR5** | **FC** | **Prionailurus** | **Faecal** |
| **83** | **201920-POR6** | **FC** | **Prionailurus** | **Faecal** |
| **84** | **201920-WPR2** | **FC** | **Prionailurus** | **Faecal** |
| **85** | **201920-WPR4** | **FC** | **Prionailurus** | **Faecal** |
| **86** | **201920-WPR5** | **FC** | **Prionailurus** | **Faecal** |
| **87** | **202223-DDW132** | **FC** | **Prionailurus** | **Faecal** |
| **88** | **202223-DDW17** | **FC** | **Prionailurus** | **Faecal** |
| **89** | **202223-DDW45** | **FC** | **Prionailurus** | **Faecal** |
| **90** | **202223-DDW47** | **FC** | **Prionailurus** | **Faecal** |
| **91** | **202223-DDW48** | **FC** | **Prionailurus** | **Faecal** |
| **92** | **202223-DDW98** | **FC** | **Prionailurus** | **Faecal** |
| **93** | **202223-PBT16** | **FC** | **Prionailurus** | **Faecal** |
| **94** | **202223-PBT161** | **FC** | **Prionailurus** | **Faecal** |
| **95** | **202223-PBT194** | **FC** | **Prionailurus** | **Faecal** |
| **96** | **CLK3** | **FC** | **Prionailurus** | **Faecal** |
| **97** | **CLK4** | **FC** | **Prionailurus** | **Faecal** |
| **98** | **PPL14** | **FC** | **Prionailurus** | **Faecal** |
| **99** | **202223-PBT262** | **LC** | **Prionailurus** | **Faecal** |
| **100** | **ANM22F28** | **LC** | **Prionailurus** | **Faecal** |
| **101** | **NAM21F07** | **LC** | **Prionailurus** | **Faecal** |
| **102** | **STR18-29** | **LC** | **Prionailurus** | **Faecal** |
| **103** | **201920-KAT2** | **RSC** | **Prionailurus** | **Faecal** |
| **104** | **201920-KIS10** | **RSC** | **Prionailurus** | **Faecal** |
| **105** | **202223-PBT246** | **RSC** | **Prionailurus** | **Faecal** |
| **106** | **I-FX-001-1** | **RSC** | **Prionailurus** | **Faecal** |

### SM4. Individual identification and population clustering

| **Table S5. Sample and marker success for each figure** | | | | | | | | |
| --- | --- | --- | --- | --- | --- | --- | --- | --- |
| **Figure** | **Total** | | **Filtering 1** | | **Filtering 2** | | | **Sample genotyping success** |
|  | **Number of sample** | **Number of marker** | **Number of sample** | **Number of marker** | **Number of sample** | **Number of marker** | **Monomorphic marker** |  |
| **2a(multispecies)** | 173 | 85 | 134 | 85 | 103 | 45 | 0 | 60% |
| **2c (Felis)** | 24 | 85 | 20 | 85 | 14 | 40 | 0 | 58% |
| **2d (Prionailurus)** | 96 | 85 | 72 | 85 | 54 | 47 | 1 | 56% |
| **2e (Panthera)** | 52 | 85 | 42 | 85 | 38 | 54 | 6 | 73% |
| **3 (Tiger)** | 29 | 85 | 26 | 85 | 19 | 55 | 14 | 66% |
| **4 (Fishing Cat)** | 75 | 85 | 57 | 85 | 40 | 36 | 4 | 53% |
| Filtering 1: Removal of poor-quality samples (replicates with >90% missing data across loci) | | | | | | | | |
| Filtering 2: Poor-efficiency markers removed (Markers with <60% sample amplification) & samples with inconclusive genotype removed (samples with <60% consensus genotype across filtered loci) | | | | | | | | |

**Table S6. Individual identification: PID and PIDsibs for species and population. *Felis* (Domestic Cat DC and Jungle Cat JC), *Prionailurus* (Fishing Cat FC, Leopard Cat LC, and Rusty-spotted Cat RSC), and *Panthera* (Tiger TG, Leopard LP, Lion LN, Snow Leopard SL).**

| **Genus** | **Species** | **Pop** | **PID** | **PIDsibs** | **Unique Individuals** |
| --- | --- | --- | --- | --- | --- |
| Felis | JC | - | 1.60E-29 | 2.80E-13 | 5 |
| Felis | DC | - | 2.50E-31 | 9.30E-14 | 9 |
| Prionailurus | LC | - | 2.60E-11 | 1.20E-05 | 4 |
| Prionailurus | RSC | - | 1.10E-16 | 3.50E-08 | 4 |
| Prionailurus | FC | AN | 1.30E-14 | 2.60E-07 | 6 |
| Prionailurus | FC | UP | 1.90E-17 | 2.00E-08 | 23 |
| Prionailurus | FC | WB | 2.60E-14 | 3.00E-07 | 3 |
| Prionailurus | FC | OD | 2.80E-10 | 2.20E-05 | 2 |
| Panthera | LN | - | 8.20E-04 | 2.80E-02 | 5 |
| Panthera | SL | - | 4.50E-08 | 2.30E-04 | 3 |
| Panthera | LP | - | 5.80E-13 | 1.70E-06 | 5 |
| Panthera | TG | Central | 3.00E-16 | 3.10E-08 | 6 |
| Panthera | TG | Northwest | 5.70E-14 | 3.00E-07 | 8 |
| Panthera | TG | Southern | 2.60E-15 | 8.10E-08 | 5 |

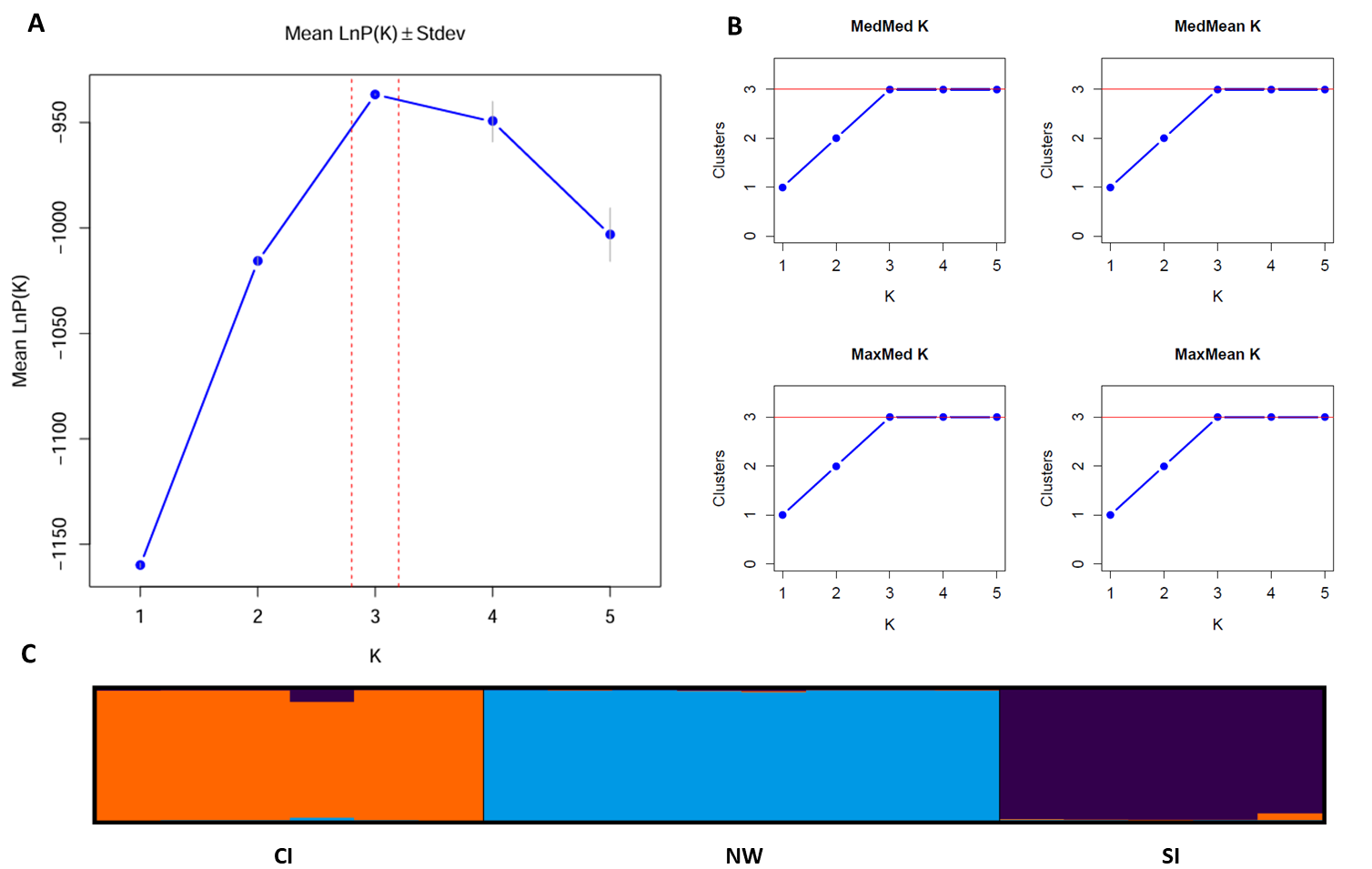

**Figure S1. Identification of genetic clusters for *Panthera tigris* samples.**

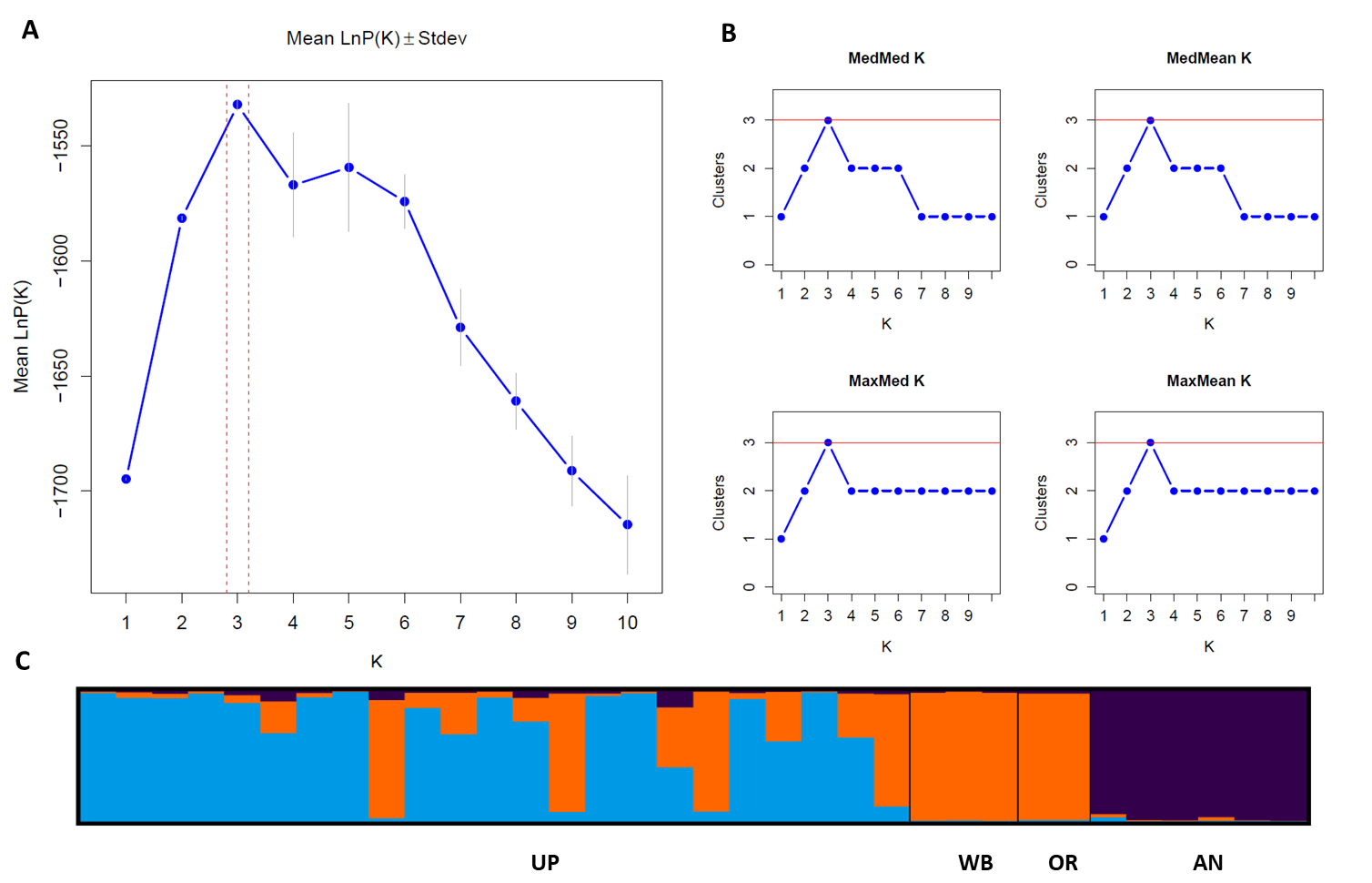

**Figure S2. Identification of genetic clusters for *Prionailurus viverrinus* (FC) samples**

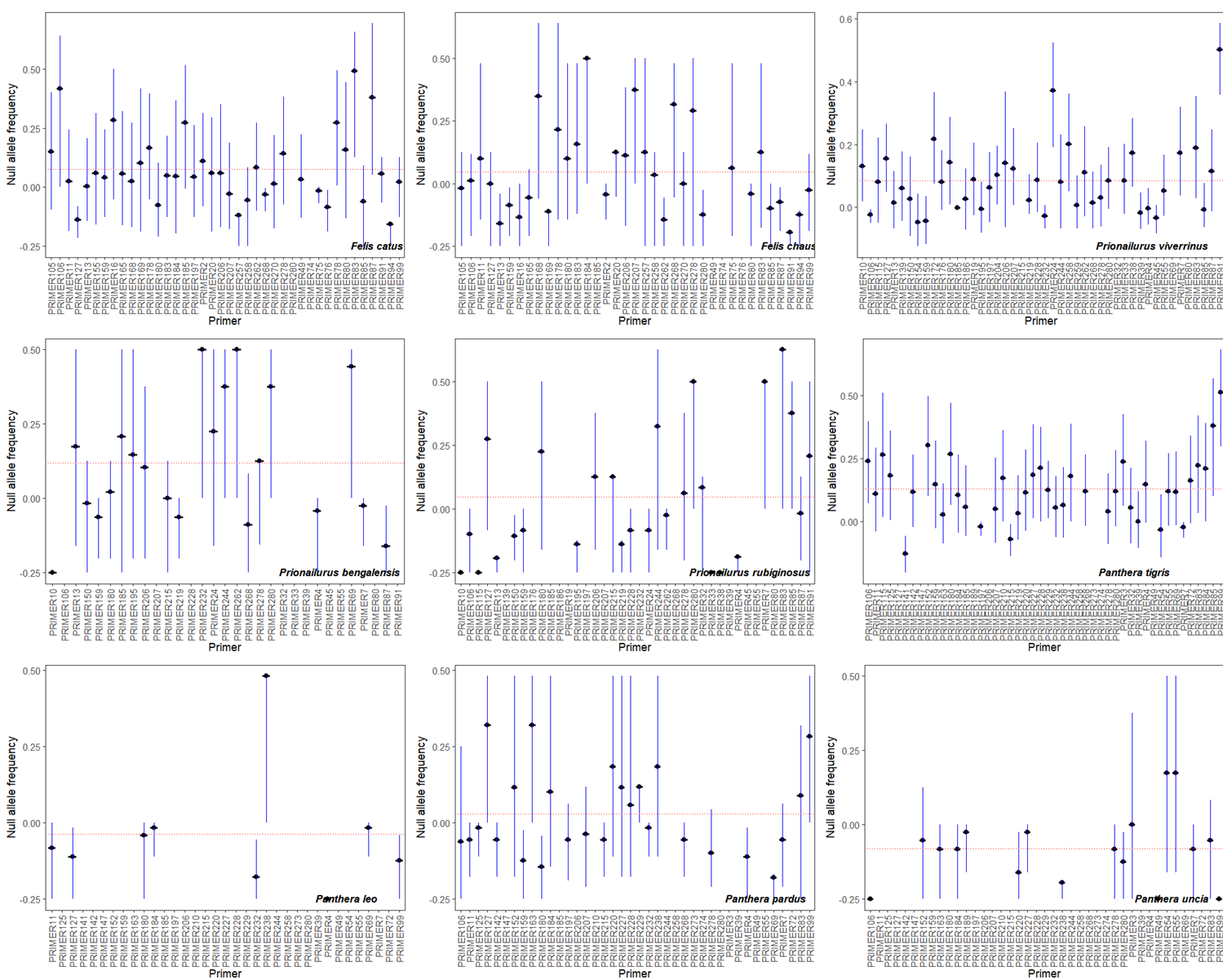

**Figure S3. Estimated per locus null allele frequencies across species. Average null allele frequencies for each species is indicated by red dotted line (Felis cactus DC = 0.075, Felis chaus JC = 0.048, Prionailurus viverrinus FC = 0.084, Prionailurus bengalensis LC = 0.119, Prionailurus rubiginosus RSC = 0.047, Panthera tigris TG = 0.13, Panthera leo LN = -0.04, Panthera pardus LP =	0.029, Panthera uncia = SL -0.08)**

| **S9. Hardy-Weinberg Equilibrium testing of markers for TG and FC** | | | | | | |
| --- | --- | --- | --- | --- | --- | --- |
| **Population** | **Marker** | **DF** | **ChiSq** | **Prob** | **Signif** | **Group** |
| **Central** | **FPX106** | **1** | **2.591** | **0.107** | **ns** | **Tiger** |
| **Central** | **FPX11** | **1** | **0.05** | **0.824** | **ns** | **Tiger** |
| **Central** | **FPX115** | **1** | **4** | **0.046** | ***** | **Tiger** |
| **Central** | **FPX125** | **1** | **5** | **0.025** | ***** | **Tiger** |
| **Central** | **FPX141** | **1** | **0.75** | **0.386** | **ns** | **Tiger** |
| **Central** | **FPX142** | **6** | **2.388** | **0.881** | **ns** | **Tiger** |
| **Central** | **FPX152** | **3** | **3.36** | **0.339** | **ns** | **Tiger** |
| **Central** | **FPX159** | **1** | **2.591** | **0.107** | **ns** | **Tiger** |
| **Central** | **FPX163** | **3** | **1.102** | **0.777** | **ns** | **Tiger** |
| **Central** | **FPX180** | **3** | **1.167** | **0.761** | **ns** | **Tiger** |
| **Central** | **FPX184** | **1** | **2.591** | **0.107** | **ns** | **Tiger** |
| **Central** | **FPX185** | **1** | **0.12** | **0.729** | **ns** | **Tiger** |
| **Central** | **FPX197** | **1** | **0.444** | **0.505** | **ns** | **Tiger** |
| **Central** | **FPX207** | **1** | **1.44** | **0.23** | **ns** | **Tiger** |
| **Central** | **FPX210** | **Monomorphic** | |  |  | **Tiger** |
| **Central** | **FPX215** | **3** | **3.467** | **0.325** | **ns** | **Tiger** |
| **Central** | **FPX219** | **Monomorphic** | |  |  | **Tiger** |
| **Central** | **FPX220** | **1** | **0.375** | **0.54** | **ns** | **Tiger** |
| **Central** | **FPX227** | **3** | **5.556** | **0.135** | **ns** | **Tiger** |
| **Central** | **FPX228** | **1** | **1.852** | **0.174** | **ns** | **Tiger** |
| **Central** | **FPX229** | **3** | **2.222** | **0.528** | **ns** | **Tiger** |
| **Central** | **FPX232** | **6** | **4** | **0.677** | **ns** | **Tiger** |
| **Central** | **FPX238** | **1** | **0.005** | **0.944** | **ns** | **Tiger** |
| **Central** | **FPX268** | **3** | **1.44** | **0.696** | **ns** | **Tiger** |
| **Central** | **FPX278** | **3** | **6.98** | **0.073** | **ns** | **Tiger** |
| **Central** | **FPX280** | **1** | **0.139** | **0.709** | **ns** | **Tiger** |
| **Central** | **FPX3** | **Monomorphic** | |  |  | **Tiger** |
| **Central** | **FPX32** | **Monomorphic** | |  |  | **Tiger** |
| **Central** | **FPX39** | **3** | **3.333** | **0.343** | **ns** | **Tiger** |
| **Central** | **FPX4** | **6** | **5.625** | **0.466** | **ns** | **Tiger** |
| **Central** | **FPX54** | **3** | **1.44** | **0.696** | **ns** | **Tiger** |
| **Central** | **FPX55** | **Monomorphic** | |  |  | **Tiger** |
| **Central** | **FPX69** | **6** | **6.5** | **0.37** | **ns** | **Tiger** |
| **Central** | **FPX7** | **1** | **0.062** | **0.804** | **ns** | **Tiger** |
| **Central** | **FPX72** | **1** | **0.667** | **0.414** | **ns** | **Tiger** |
| **Central** | **FPX75** | **Monomorphic** | |  |  | **Tiger** |
| **Central** | **FPX83** | **1** | **0.444** | **0.505** | **ns** | **Tiger** |
| **Central** | **FPX84** | **Monomorphic** | |  |  | **Tiger** |
| **Central** | **FPX85** | **1** | **2** | **0.157** | **ns** | **Tiger** |
| **Central** | **FPX99** | **1** | **0.139** | **0.709** | **ns** | **Tiger** |
| **Northwest** | **FPX106** | **1** | **0.163** | **0.686** | **ns** | **Tiger** |
| **Northwest** | **FPX11** | **1** | **2.16** | **0.142** | **ns** | **Tiger** |
| **Northwest** | **FPX115** | **1** | **0.2** | **0.655** | **ns** | **Tiger** |
| **Northwest** | **FPX125** | **Monomorphic** | |  |  | **Tiger** |
| **Northwest** | **FPX141** | **1** | **2.222** | **0.136** | **ns** | **Tiger** |
| **Northwest** | **FPX142** | **3** | **7.147** | **0.067** | **ns** | **Tiger** |
| **Northwest** | **FPX152** | **3** | **1.44** | **0.696** | **ns** | **Tiger** |
| **Northwest** | **FPX159** | **Monomorphic** | |  |  | **Tiger** |
| **Northwest** | **FPX163** | **1** | **0.667** | **0.414** | **ns** | **Tiger** |
| **Northwest** | **FPX180** | **1** | **0.163** | **0.686** | **ns** | **Tiger** |
| **Northwest** | **FPX184** | **1** | **2.88** | **0.09** | **ns** | **Tiger** |
| **Northwest** | **FPX185** | **Monomorphic** | |  |  | **Tiger** |
| **Northwest** | **FPX197** | **1** | **0.062** | **0.804** | **ns** | **Tiger** |
| **Northwest** | **FPX207** | **1** | **0.313** | **0.576** | **ns** | **Tiger** |
| **Northwest** | **FPX210** | **1** | **4** | **0.046** | ***** | **Tiger** |
| **Northwest** | **FPX215** | **1** | **0.889** | **0.346** | **ns** | **Tiger** |
| **Northwest** | **FPX219** | **1** | **0.036** | **0.85** | **ns** | **Tiger** |
| **Northwest** | **FPX220** | **1** | **0.889** | **0.346** | **ns** | **Tiger** |
| **Northwest** | **FPX227** | **3** | **1.05** | **0.789** | **ns** | **Tiger** |
| **Northwest** | **FPX228** | **Monomorphic** | |  |  | **Tiger** |
| **Northwest** | **FPX229** | **1** | **0.75** | **0.386** | **ns** | **Tiger** |
| **Northwest** | **FPX232** | **3** | **4.212** | **0.239** | **ns** | **Tiger** |
| **Northwest** | **FPX238** | **1** | **0.583** | **0.445** | **ns** | **Tiger** |
| **Northwest** | **FPX268** | **3** | **0.918** | **0.821** | **ns** | **Tiger** |
| **Northwest** | **FPX278** | **1** | **4.84** | **0.028** | ***** | **Tiger** |
| **Northwest** | **FPX280** | **1** | **0.24** | **0.624** | **ns** | **Tiger** |
| **Northwest** | **FPX3** | **3** | **8.16** | **0.043** | ***** | **Tiger** |
| **Northwest** | **FPX32** | **1** | **0.444** | **0.505** | **ns** | **Tiger** |
| **Northwest** | **FPX39** | **3** | **2.062** | **0.56** | **ns** | **Tiger** |
| **Northwest** | **FPX4** | **3** | **4.587** | **0.205** | **ns** | **Tiger** |
| **Northwest** | **FPX54** | **3** | **5.556** | **0.135** | **ns** | **Tiger** |
| **Northwest** | **FPX55** | **1** | **0.454** | **0.501** | **ns** | **Tiger** |
| **Northwest** | **FPX69** | **1** | **2** | **0.157** | **ns** | **Tiger** |
| **Northwest** | **FPX7** | **Monomorphic** | |  |  | **Tiger** |
| **Northwest** | **FPX72** | **Monomorphic** | |  |  | **Tiger** |
| **Northwest** | **FPX75** | **Monomorphic** | |  |  | **Tiger** |
| **Northwest** | **FPX83** | **1** | **0.375** | **0.54** | **ns** | **Tiger** |
| **Northwest** | **FPX84** | **Monomorphic** | |  |  | **Tiger** |
| **Northwest** | **FPX85** | **Monomorphic** | |  |  | **Tiger** |
| **Northwest** | **FPX99** | **1** | **0.194** | **0.659** | **ns** | **Tiger** |
| **Southern** | **FPX106** | **Monomorphic** | |  |  | **Tiger** |
| **Southern** | **FPX11** | **3** | **5** | **0.172** | **ns** | **Tiger** |
| **Southern** | **FPX115** | **3** | **6** | **0.112** | **ns** | **Tiger** |
| **Southern** | **FPX125** | **1** | **0** | **1** | **ns** | **Tiger** |
| **Southern** | **FPX141** | **1** | **4** | **0.046** | ***** | **Tiger** |
| **Southern** | **FPX142** | **6** | **1.44** | **0.963** | **ns** | **Tiger** |
| **Southern** | **FPX152** | **1** | **0** | **1** | **ns** | **Tiger** |
| **Southern** | **FPX159** | **1** | **0.918** | **0.338** | **ns** | **Tiger** |
| **Southern** | **FPX163** | **3** | **2.778** | **0.427** | **ns** | **Tiger** |
| **Southern** | **FPX180** | **1** | **4** | **0.046** | ***** | **Tiger** |
| **Southern** | **FPX184** | **3** | **8** | **0.046** | ***** | **Tiger** |
| **Southern** | **FPX185** | **1** | **0.139** | **0.709** | **ns** | **Tiger** |
| **Southern** | **FPX197** | **1** | **0.082** | **0.775** | **ns** | **Tiger** |
| **Southern** | **FPX207** | **1** | **0.082** | **0.775** | **ns** | **Tiger** |
| **Southern** | **FPX210** | **Monomorphic** | |  |  | **Tiger** |
| **Southern** | **FPX215** | **1** | **0.313** | **0.576** | **ns** | **Tiger** |
| **Southern** | **FPX219** | **1** | **0.313** | **0.576** | **ns** | **Tiger** |
| **Southern** | **FPX220** | **3** | **3.36** | **0.339** | **ns** | **Tiger** |
| **Southern** | **FPX227** | **1** | **0.918** | **0.338** | **ns** | **Tiger** |
| **Southern** | **FPX228** | **1** | **5** | **0.025** | ***** | **Tiger** |
| **Southern** | **FPX229** | **Monomorphic** | |  |  | **Tiger** |
| **Southern** | **FPX232** | **1** | **0.871** | **0.351** | **ns** | **Tiger** |
| **Southern** | **FPX238** | **1** | **4** | **0.046** | ***** | **Tiger** |
| **Southern** | **FPX268** | **3** | **1.444** | **0.695** | **ns** | **Tiger** |
| **Southern** | **FPX278** | **1** | **0.918** | **0.338** | **ns** | **Tiger** |
| **Southern** | **FPX280** | **1** | **0.918** | **0.338** | **ns** | **Tiger** |
| **Southern** | **FPX3** | **1** | **0.444** | **0.505** | **ns** | **Tiger** |
| **Southern** | **FPX32** | **3** | **1.44** | **0.696** | **ns** | **Tiger** |
| **Southern** | **FPX39** | **6** | **4** | **0.677** | **ns** | **Tiger** |
| **Southern** | **FPX4** | **3** | **5** | **0.172** | **ns** | **Tiger** |
| **Southern** | **FPX54** | **1** | **0.444** | **0.505** | **ns** | **Tiger** |
| **Southern** | **FPX55** | **Monomorphic** | |  |  | **Tiger** |
| **Southern** | **FPX69** | **1** | **0.139** | **0.709** | **ns** | **Tiger** |
| **Southern** | **FPX7** | **1** | **0.313** | **0.576** | **ns** | **Tiger** |
| **Southern** | **FPX72** | **1** | **0.062** | **0.804** | **ns** | **Tiger** |
| **Southern** | **FPX75** | **Monomorphic** | |  |  | **Tiger** |
| **Southern** | **FPX83** | **1** | **0.062** | **0.804** | **ns** | **Tiger** |
| **Southern** | **FPX84** | **1** | **0** | **1** | **ns** | **Tiger** |
| **Southern** | **FPX85** | **Monomorphic** | |  |  | **Tiger** |
| **Southern** | **FPX99** | **1** | **4** | **0.046** | ***** | **Tiger** |
| **AN** | **FPX10** | **1** | **0.24** | **0.624** | **ns** | **Fishing Cat** |
| **AN** | **FPX106** | **1** | **0.24** | **0.624** | **ns** | **Fishing Cat** |
| **AN** | **FPX127** | **3** | **7.28** | **0.063** | **ns** | **Fishing Cat** |
| **AN** | **FPX13** | **3** | **2.987** | **0.394** | **ns** | **Fishing Cat** |
| **AN** | **FPX139** | **3** | **2.333** | **0.506** | **ns** | **Fishing Cat** |
| **AN** | **FPX154** | **3** | **4** | **0.261** | **ns** | **Fishing Cat** |
| **AN** | **FPX159** | **1** | **0.667** | **0.414** | **ns** | **Fishing Cat** |
| **AN** | **FPX172** | **1** | **4** | **0.046** | ***** | **Fishing Cat** |
| **AN** | **FPX176** | **Monomorphic** | |  |  | **Fishing Cat** |
| **AN** | **FPX185** | **Monomorphic** | |  |  | **Fishing Cat** |
| **AN** | **FPX186** | **6** | **14.28** | **0.027** | ***** | **Fishing Cat** |
| **AN** | **FPX19** | **1** | **0.12** | **0.729** | **ns** | **Fishing Cat** |
| **AN** | **FPX195** | **6** | **3.889** | **0.692** | **ns** | **Fishing Cat** |
| **AN** | **FPX197** | **3** | **2.45** | **0.484** | **ns** | **Fishing Cat** |
| **AN** | **FPX204** | **1** | **4** | **0.046** | ***** | **Fishing Cat** |
| **AN** | **FPX207** | **3** | **2.8** | **0.423** | **ns** | **Fishing Cat** |
| **AN** | **FPX219** | **1** | **0.005** | **0.944** | **ns** | **Fishing Cat** |
| **AN** | **FPX228** | **3** | **7.778** | **0.051** | **ns** | **Fishing Cat** |
| **AN** | **FPX232** | **1** | **0.313** | **0.576** | **ns** | **Fishing Cat** |
| **AN** | **FPX258** | **6** | **5** | **0.544** | **ns** | **Fishing Cat** |
| **AN** | **FPX259** | **1** | **4** | **0.046** | ***** | **Fishing Cat** |
| **AN** | **FPX268** | **1** | **0.082** | **0.775** | **ns** | **Fishing Cat** |
| **AN** | **FPX278** | **1** | **0.521** | **0.471** | **ns** | **Fishing Cat** |
| **AN** | **FPX280** | **3** | **1.73** | **0.63** | **ns** | **Fishing Cat** |
| **AN** | **FPX33** | **1** | **2.16** | **0.142** | **ns** | **Fishing Cat** |
| **AN** | **FPX38** | **1** | **0** | **1** | **ns** | **Fishing Cat** |
| **AN** | **FPX39** | **1** | **0.194** | **0.659** | **ns** | **Fishing Cat** |
| **AN** | **FPX4** | **3** | **1.389** | **0.708** | **ns** | **Fishing Cat** |
| **AN** | **FPX55** | **1** | **0.24** | **0.624** | **ns** | **Fishing Cat** |
| **AN** | **FPX7** | **6** | **6.25** | **0.396** | **ns** | **Fishing Cat** |
| **AN** | **FPX85** | **1** | **0.313** | **0.576** | **ns** | **Fishing Cat** |
| **AN** | **FPX91** | **3** | **3.333** | **0.343** | **ns** | **Fishing Cat** |
| **UP** | **FPX10** | **1** | **13.405** | **0** | ******* | **Fishing Cat** |
| **UP** | **FPX106** | **6** | **2.204** | **0.9** | **ns** | **Fishing Cat** |
| **UP** | **FPX127** | **6** | **21.344** | **0.002** | ****** | **Fishing Cat** |
| **UP** | **FPX13** | **3** | **0.985** | **0.805** | **ns** | **Fishing Cat** |
| **UP** | **FPX139** | **3** | **1.37** | **0.713** | **ns** | **Fishing Cat** |
| **UP** | **FPX154** | **6** | **1.971** | **0.922** | **ns** | **Fishing Cat** |
| **UP** | **FPX159** | **3** | **3.783** | **0.286** | **ns** | **Fishing Cat** |
| **UP** | **FPX172** | **10** | **22.401** | **0.013** | ***** | **Fishing Cat** |
| **UP** | **FPX176** | **1** | **4.973** | **0.026** | ***** | **Fishing Cat** |
| **UP** | **FPX185** | **1** | **0.04** | **0.842** | **ns** | **Fishing Cat** |
| **UP** | **FPX186** | **6** | **1.356** | **0.968** | **ns** | **Fishing Cat** |
| **UP** | **FPX19** | **6** | **22.447** | **0.001** | ****** | **Fishing Cat** |
| **UP** | **FPX195** | **6** | **5.564** | **0.474** | **ns** | **Fishing Cat** |
| **UP** | **FPX197** | **3** | **4.493** | **0.213** | **ns** | **Fishing Cat** |
| **UP** | **FPX204** | **3** | **0.893** | **0.827** | **ns** | **Fishing Cat** |
| **UP** | **FPX207** | **6** | **15.531** | **0.017** | ***** | **Fishing Cat** |
| **UP** | **FPX219** | **3** | **0.281** | **0.964** | **ns** | **Fishing Cat** |
| **UP** | **FPX228** | **3** | **0.771** | **0.856** | **ns** | **Fishing Cat** |
| **UP** | **FPX232** | **3** | **0.86** | **0.835** | **ns** | **Fishing Cat** |
| **UP** | **FPX258** | **15** | **33.639** | **0.004** | ****** | **Fishing Cat** |
| **UP** | **FPX259** | **1** | **0.007** | **0.933** | **ns** | **Fishing Cat** |
| **UP** | **FPX268** | **3** | **0.7** | **0.873** | **ns** | **Fishing Cat** |
| **UP** | **FPX278** | **6** | **3.133** | **0.792** | **ns** | **Fishing Cat** |
| **UP** | **FPX280** | **6** | **12.666** | **0.049** | ***** | **Fishing Cat** |
| **UP** | **FPX33** | **3** | **2.39** | **0.496** | **ns** | **Fishing Cat** |
| **UP** | **FPX38** | **6** | **31.466** | **0** | ******* | **Fishing Cat** |
| **UP** | **FPX39** | **6** | **3.67** | **0.721** | **ns** | **Fishing Cat** |
| **UP** | **FPX4** | **6** | **2.865** | **0.826** | **ns** | **Fishing Cat** |
| **UP** | **FPX55** | **1** | **3.185** | **0.074** | **ns** | **Fishing Cat** |
| **UP** | **FPX7** | **6** | **13.267** | **0.039** | ***** | **Fishing Cat** |
| **UP** | **FPX85** | **3** | **0.301** | **0.96** | **ns** | **Fishing Cat** |
| **UP** | **FPX91** | **3** | **30.661** | **0** | ******* | **Fishing Cat** |
| **WB** | **FPX10** | **1** | **4** | **0.046** | ***** | **Fishing Cat** |
| **WB** | **FPX106** | **1** | **0.444** | **0.505** | **ns** | **Fishing Cat** |
| **WB** | **FPX127** | **6** | **6** | **0.423** | **ns** | **Fishing Cat** |
| **WB** | **FPX13** | **3** | **2.333** | **0.506** | **ns** | **Fishing Cat** |
| **WB** | **FPX139** | **3** | **2.333** | **0.506** | **ns** | **Fishing Cat** |
| **WB** | **FPX154** | **1** | **0.222** | **0.637** | **ns** | **Fishing Cat** |
| **WB** | **FPX159** | **1** | **0.75** | **0.386** | **ns** | **Fishing Cat** |
| **WB** | **FPX172** | **1** | **2** | **0.157** | **ns** | **Fishing Cat** |
| **WB** | **FPX176** | **1** | **0.082** | **0.775** | **ns** | **Fishing Cat** |
| **WB** | **FPX185** | **Monomorphic** | |  |  | **Fishing Cat** |
| **WB** | **FPX186** | **1** | **0** | **1** | **ns** | **Fishing Cat** |
| **WB** | **FPX19** | **3** | **2** | **0.572** | **ns** | **Fishing Cat** |
| **WB** | **FPX195** | **3** | **3.333** | **0.343** | **ns** | **Fishing Cat** |
| **WB** | **FPX197** | **3** | **6** | **0.112** | **ns** | **Fishing Cat** |
| **WB** | **FPX204** | **1** | **0.222** | **0.637** | **ns** | **Fishing Cat** |
| **WB** | **FPX207** | **6** | **4.889** | **0.558** | **ns** | **Fishing Cat** |
| **WB** | **FPX219** | **1** | **0.12** | **0.729** | **ns** | **Fishing Cat** |
| **WB** | **FPX228** | **3** | **1.444** | **0.695** | **ns** | **Fishing Cat** |
| **WB** | **FPX232** | **1** | **0.444** | **0.505** | **ns** | **Fishing Cat** |
| **WB** | **FPX258** | **Monomorphic** | |  |  | **Fishing Cat** |
| **WB** | **FPX259** | **1** | **0.222** | **0.637** | **ns** | **Fishing Cat** |
| **WB** | **FPX268** | **1** | **4** | **0.046** | ***** | **Fishing Cat** |
| **WB** | **FPX278** | **1** | **1.44** | **0.23** | **ns** | **Fishing Cat** |
| **WB** | **FPX280** | **1** | **0.12** | **0.729** | **ns** | **Fishing Cat** |
| **WB** | **FPX33** | **1** | **0.871** | **0.351** | **ns** | **Fishing Cat** |
| **WB** | **FPX38** | **Monomorphic** | |  |  | **Fishing Cat** |
| **WB** | **FPX39** | **1** | **0.444** | **0.505** | **ns** | **Fishing Cat** |
| **WB** | **FPX4** | **3** | **4.16** | **0.245** | **ns** | **Fishing Cat** |
| **WB** | **FPX55** | **1** | **0.082** | **0.775** | **ns** | **Fishing Cat** |
| **WB** | **FPX7** | **3** | **3.333** | **0.343** | **ns** | **Fishing Cat** |
| **WB** | **FPX85** | **3** | **3** | **0.392** | **ns** | **Fishing Cat** |
| **WB** | **FPX91** | **3** | **4.444** | **0.217** | **ns** | **Fishing Cat** |
| **OD** | **FPX10** | **Monomorphic** | |  |  | **Fishing Cat** |
| **OD** | **FPX106** | **Monomorphic** | |  |  | **Fishing Cat** |
| **OD** | **FPX127** | **Monomorphic** | |  |  | **Fishing Cat** |
| **OD** | **FPX13** | **3** | **2** | **0.572** | **ns** | **Fishing Cat** |
| **OD** | **FPX139** | **Monomorphic** | |  |  | **Fishing Cat** |
| **OD** | **FPX154** | **3** | **2** | **0.572** | **ns** | **Fishing Cat** |
| **OD** | **FPX159** | **3** | **2** | **0.572** | **ns** | **Fishing Cat** |
| **OD** | **FPX172** | **3** | **2** | **0.572** | **ns** | **Fishing Cat** |
| **OD** | **FPX176** | **Monomorphic** | |  |  | **Fishing Cat** |
| **OD** | **FPX185** | **Monomorphic** | |  |  | **Fishing Cat** |
| **OD** | **FPX186** | **1** | **2** | **0.157** | **ns** | **Fishing Cat** |
| **OD** | **FPX19** | **1** | **2** | **0.157** | **ns** | **Fishing Cat** |
| **OD** | **FPX195** | **1** | **0.222** | **0.637** | **ns** | **Fishing Cat** |
| **OD** | **FPX197** | **1** | **0.222** | **0.637** | **ns** | **Fishing Cat** |
| **OD** | **FPX204** | **Monomorphic** | |  |  | **Fishing Cat** |
| **OD** | **FPX207** | **1** | **0.222** | **0.637** | **ns** | **Fishing Cat** |
| **OD** | **FPX219** | **Monomorphic** | |  |  | **Fishing Cat** |
| **OD** | **FPX228** | **1** | **2** | **0.157** | **ns** | **Fishing Cat** |
| **OD** | **FPX232** | **1** | **0.222** | **0.637** | **ns** | **Fishing Cat** |
| **OD** | **FPX258** | **3** | **2** | **0.572** | **ns** | **Fishing Cat** |
| **OD** | **FPX259** | **Monomorphic** | |  |  | **Fishing Cat** |
| **OD** | **FPX268** | **1** | **0.222** | **0.637** | **ns** | **Fishing Cat** |
| **OD** | **FPX278** | **Monomorphic** | |  |  | **Fishing Cat** |
| **OD** | **FPX280** | **3** | **2** | **0.572** | **ns** | **Fishing Cat** |
| **OD** | **FPX33** | **1** | **0.222** | **0.637** | **ns** | **Fishing Cat** |
| **OD** | **FPX38** | **1** | **0.222** | **0.637** | **ns** | **Fishing Cat** |
| **OD** | **FPX39** | **Monomorphic** | |  |  | **Fishing Cat** |
| **OD** | **FPX4** | **Monomorphic** | |  |  | **Fishing Cat** |
| **OD** | **FPX55** | **1** | **0.222** | **0.637** | **ns** | **Fishing Cat** |
| **OD** | **FPX7** | **3** | **4** | **0.261** | **ns** | **Fishing Cat** |
| **OD** | **FPX85** | **1** | **1** | **0.317** | **ns** | **Fishing Cat** |
| **OD** | **FPX91** | **1** | **2** | **0.157** | **ns** | **Fishing Cat** |

| **Table S10. Genotyping error rates per loci for different types of samples** | | | | | |
| --- | --- | --- | --- | --- | --- |
| **Invasive samples** | | | **Non-invasive samples** | | |
| Loci | Allelic Dropout rate | False allele rate | Loci | Allelic Dropout rate | False allele rate |
| FPX106 | 0 | 0 | FPX10 | 0 | 0 |
| FPX11 | 0 | 0.017544 | FPX106 | 0 | 0 |
| FPX115 | 0 | 0 | FPX127 | 0 | 0 |
| FPX125 | 0 | 0 | FPX13 | 0 | 0 |
| FPX127 | 0 | 0 | FPX139 | 0 | 0 |
| FPX13 | 0 | 0.025641 | FPX154 | 0 | 0 |
| FPX141 | 0.090909 | 0 | FPX159 | 0 | 0 |
| FPX142 | 0.027778 | 0 | FPX172 | 0.019231 | 0 |
| FPX147 | 0 | 0 | FPX176 | 0 | 0 |
| FPX152 | 0.095238 | 0 | FPX185 | 0 | 0 |
| FPX159 | 0.111111 | 0 | FPX186 | 0.043182 | 0 |
| FPX163 | 0 | 0 | FPX19 | 0.03 | 0 |
| FPX172 | 0 | 0 | FPX195 | 0.068667 | 0.007143 |
| FPX180 | 0 | 0.015873 | FPX197 | 0 | 0 |
| FPX184 | 0 | 0.015873 | FPX204 | 0 | 0 |
| FPX185 | 0 | 0.222222 | FPX207 | 0.066667 | 0.027778 |
| FPX189 | 0 | 0 | FPX215 | 0 | 0 |
| FPX19 | 0 | 0 | FPX219 | 0 | 0 |
| FPX197 | 0 | 0 | FPX228 | 0.068254 | 0 |
| FPX204 | 0 | 0 | FPX232 | 0 | 0 |
| FPX206 | 0 | 0 | FPX258 | 0.02 | 0.042754 |
| FPX207 | 0 | 0 | FPX259 | 0 | 0 |
| FPX210 | 0 | 0.019608 | FPX268 | 0 | 0 |
| FPX215 | 0 | 0 | FPX278 | 0.011905 | 0 |
| FPX219 | 0 | 0 | FPX280 | 0 | 0 |
| FPX220 | 0 | 0 | FPX32 | 0 | 0 |
| FPX227 | 0.047619 | 0.039216 | FPX33 | 0 | 0 |
| FPX228 | 0 | 0 | FPX38 | 0 | 0 |
| FPX229 | 0 | 0 | FPX39 | 0 | 0 |
| FPX232 | 0 | 0 | FPX4 | 0 | 0 |
| FPX238 | 0 | 0 | FPX55 | 0.153333 | 0.008772 |
| FPX244 | 0 | 0 | FPX7 | 0.103509 | 0.009259 |
| FPX258 | 0 | 0 | FPX76 | 0 | 0 |
| FPX259 | 0 | 0 | FPX80 | 0 | 0 |
| FPX268 | 0.037037 | 0 | FPX85 | 0.02381 | 0.007143 |
| FPX273 | 0 | 0 | FPX91 | 0.281905 | 0.042342 |
| FPX274 | 0 | 0 |  |  |  |
| FPX278 | 0 | 0.015152 |  |  |  |
| FPX280 | 0 | 0 |  |  |  |
| FPX3 | 0 | 0 |  |  |  |
| FPX32 | 0 | 0 |  |  |  |
| FPX39 | 0 | 0 |  |  |  |
| FPX4 | 0 | 0 |  |  |  |
| FPX49 | 0 | 0.018519 |  |  |  |
| FPX54 | 0 | 0 |  |  |  |
| FPX55 | 0 | 0 |  |  |  |
| FPX69 | 0.030303 | 0 |  |  |  |
| FPX7 | 0 | 0 |  |  |  |
| FPX72 | 0 | 0 |  |  |  |
| FPX75 | 0 | 0.055556 |  |  |  |
| FPX76 | 0 | 0 |  |  |  |
| FPX83 | 0 | 0 |  |  |  |
| FPX84 | 0.166667 | 0 |  |  |  |
| FPX85 | 0 | 0 |  |  |  |
| FPX99 | 0 | 0.018519 |  |  |  |
